# RIG-I activation primes and trains innate antiviral immune memory

**DOI:** 10.1101/2022.10.27.514004

**Authors:** Maike S Adamson, Svetozar Nesic, Andreas Buness, Kübra Bayrak, Saskia Schmitz, Sofía Soler, Thomas Zillinger, Samira Marx, Silke Lambing, Katarzyna Andryka-Cegielski, Fawad Khan, Thais M. Schlee-Guimarães, Stephan Herberhold, Michele Proietti, Katarzyna Placek, Patrick Reading, Luca Cicin-Sain, Martin Schlee, Jasper van den Boorn, Evelyn Hartmann, Gunther Hartmann, Eva Bartok

**Affiliations:** Institute of Clinical Chemistry and Clinical Pharmacology, University Hospital Bonn, Bonn, Germany; Institute of Experimental Haematology and Transfusion Medicine, University Hospital, Bonn, Germany; Core Unit for Bioinformatics Data Analysis, Institute of Medical Biometry, Informatics and Epidemiology, Bonn, Germany; Helmholtz Centre for Infection Research, Department of Viral Immunology, Braunschweig, Germany; Department of Otorhinolaryngology, Johanniter Waldkrankenhaus, Bonn, Germany; Center for Chronic Immunodeficiency, University Hospital Freiburg, Germany; Life and Medical Sciences Institute, University of Bonn, Germany; The Peter Doherty Institute for Infection and Immunity, Melbourne, Victoria, Australia; German Center for Infection Research (DZIF), Braunschweig, Germany; Department of Otorhinolaryngology, University Hospital Bonn, 53127 Bonn, Germany; Unit of Experimental Immunology, Institute of Tropical Medicine, Antwerp, Belgium

## Abstract

Adaptive processes of the innate immune system, known as trained immunity (TI), are critical to human health and disease, yet they have not been systematically investigated downstream of antiviral sensing. Here, we elucidate the potential of the antiviral cytosolic RNA receptor retinoic acid-inducible gene I (RIG-I) to train, prime and tolerize the innate immune system. Using a specific RIG-I agonist, we observed that repetitive stimulation enhanced interferon-stimulated gene (ISG) and pro-inflammatory cytokine induction in human primary monocytes, epithelial cells and fibroblasts and afforded non-specific antiviral protection. RNA sequencing revealed broad, cell type-specific transcriptional changes, indicative of priming of ISGs and training of the NFκB pathway, without measurable tolerization, while ATAC sequencing in monocytes demonstrated chromatin remodeling and enhanced accessibility of key transcription factor-binding motifs such as STAT1. Moreover, while STAT1 signaling was critically required, it was not sufficient to recapitulate RIG-I induced TI. Altogether, our data demonstrate that RIG-I-mediated TI promotes an immunologically alert state with important implications for host defense and the application of RIG-I ligands in anti-infective and anti-tumoral therapies.

**One Sentence Summary:** RIG-I activation trains and primes innate immune response at the cellular level, affording non-specific immune protection by immune and non-immune cells.

## INTRODUCTION

Numerous recent publications have underscored the importance of innate immune training and memory for human health and disease(Netea *et al*, 2020; Berthelot & Sibilia, 2019; Mantovani & Netea, 2020). Immune training, along with priming and tolerization, all refer to types of long-term functional reprogramming of the innate immune response(Netea *et al*, 2020; Divangahi *et al*, 2021a). While tolerization denotes a reduction in response upon restimulation, as has long been described for LPS and other endotoxins(Freudenberg & Galanos, 1988), priming and training both describe differential states of heightened reactivity. Priming denotes long-term changes in the immune status, such as long-term transcription of interferon-stimulated genes (ISG) after type-I interferon (IFN) exposure(Kamada *et al*, 2018), while, in contrast, training describes a state of increased responsiveness upon restimulation despite the lack of a prior, residual “priming” signal. Innate immune training has been reported and characterized for multiple immune stimuli(Netea *et al*, 2020), including most prominently the fungal wall component and Dectin-1 ligand *β*-glucan(Quintin *et al*, 2012) and the *Mycobacterium* vaccine strain, Bacille Calmette-Guérin (BCG)(Kleinnijenhuis *et al*, 2012; Arts *et al*, 2018b). *β*-glucan exposure has been shown to promote responses to endotoxins and even reverse the tolerance induced by LPS (Quintin *et al*, 2012; Novakovic *et al*, 2016). BCG has been known for decades to induce heterologous immune protection, including resistance to infection with babesia and plasmodia *spp*., influenza A virus (IAV) and herpes simplex 2 virus in mice(Clark *et al*, 1976; Spencer *et al*, 1977; Starr *et al*, 1976), antitumoral effects during bladder cancer(Morales *et al*, 1976), and a non-specific reduction of mortality among vaccinated children(Näslund; Faustman & Davis, 2015). In recent years, the molecular mechanisms underlying both *β*-glucan-and BCG vaccination-mediated training have been explored systematically *in vivo* and in standardized *in vitro* protocols with primary human monocytes(Kleinnijenhuis *et al*, 2014; Bekkering *et al*, 2016a; Arts *et al*, 2018b; Cirovic *et al*, 2020; Domínguez-Andrés *et al*, 2021), demonstrating that these effects are governed by metabolic reprogramming and long-term epigenetic changes in chromatin modifications and accessibility(Netea *et al*, 2020; Fanucchi *et al*, 2020; Saeed *et al*, 2014; Kleinnijenhuis *et al*, 2012). Similar findings have also been reported for diverse infectious agents, including human-pathogenic bacteria(Guillon *et al*, 2020), live vaccines(Gyssens & Netea, 2019) and plasmodia(Schrum *et al*, 2018; Walk *et al*, 2020) and are hypothesized to contribute to autoinflammatory and autoimmune disease(Arts *et al*, 2018a; Berthelot & Sibilia, 2019).

Although it has likewise been known for decades that the enzymatically-generated double-stranded (ds) RNA molecule poly(I:C) confers lasting, broad antiviral effects(Stephen *et al*, 1977; Richmond & Hamilton, 1969), only a limited number of studies have investigated innate immune training induced by nucleic-acid (NA) sensing. To date, these have focused on endosomal Toll-like receptor (TLR) activation (TLR3, 7-9) by NA(Butcher *et al*, 2018; Ifrim *et al*, 2014), and none has explored the effect of cytosolic NA sensing in this context. Intriguingly, endosomal TLR activation with NA has been shown to potentially induce both tolerization and priming of subsequent responses, depending on the DNA or RNA dose used(Ifrim *et al*, 2014). However, these observations are not transferrable to cytosolic sensing, given that their immunological functions are fundamentally different: while the endosomal TLRs are primarily expressed in immune and barrier cells and are important to sensing pathogens in the host at large, cytosolic NA sensors are broadly expressed in many cell types and act to recognize viral replication within the cell itself(Hartmann, 2017; Bartok & Hartmann, 2020).

The cytosolic RNA receptor retinoic acid-inducible gene-I (RIG-I) is expressed in virtually all nucleated cells, where it acts as an essential sensor of RNA viruses and induces type I and type III IFN signaling, NFκB activation and the expression of pro-inflammatory genes, as well as the direct induction of antiviral effectors(Schlee, 2013; Rehwinkel & Gack, 2020). Moreover, fulminant RIG-I activation can also induce proinflammatory cell death, particularly in tumor cells(van den Boorn & Hartmann, 2013). RIG-I is activated by blunt-ended dsRNA with incomplete cap methylation (cap0) or a 5’ terminal di- or triphosphate, such as found on nascent viral transcripts as well as their molecular mimic poly(I:C)(Rehwinkel & Gack, 2020). However, while poly(I:C) and other long dsRNA are prototypical danger signals which are sensed by multiple RNA receptors(Bartok & Hartmann, 2020), specific agonists for RIG-I have been developed(Junt & Barchet, 2015; Feld & Schuberth-Wagner, 2022) and are under research and development for applications in tumor immunotherapy and the prophylaxis and treatment of viral infections(Coch *et al*, 2017a; Mao *et al*, 2021; Marx *et al*, 2022; U.S.N.L. of a study evaluating the safety, pharmacokinetics, and antiviral efficacy of SB 9200 in subjects infected with chronic HBV, 2020; MK-4621-002, 2022). Thus, immune memory effects induced by the respective NA ligands could potentially affect the clinical dosage and effectivity in the context of prophylactic and therapeutic applications. Moreover, the broad expression of RIG-I in both immune and non-immune cells means that these effects could potentially occur in almost any cell and even differ between cell types.

In the current study, we investigated potential adaptive changes in the innate immune response to repetitive activation (day 0 and/or day 6) of RIG-I in human primary monocytes, fibroblasts, and epithelial cells, following established immune training protocols(Bekkering *et al*, 2016a; Domínguez-Andrés *et al*, 2021). RIG-I activation induced broad, transcriptional changes in all three cell types, demonstrating immune priming of antiviral genes and training, particularly of NFκB signaling, without any substantial tolerization. Moreover, RIG-I-induced trained immunity (TI) protected cells non-specifically against both the RNA virus influenza A and the DNA virus CMV. While all three cell types demonstrated priming and training of immune-related genes, key cell-cycle genes were also differentially regulated in monocytes but not in fibroblasts and epithelial cells, indicating an important difference in the response of myeloid and longer-lived non-immune cells to RIG-I activation. Moreover, pathway analysis after RIG-I-training revealed long-term upregulation of multiple innate immune sensors and downstream signaling molecules, including an unexpected sensitization for the P2X7/NLRP3 inflammasome pathway, which likely contributes to the antiviral and antitumoral effects of RIG-I stimulation. At the epigenetic level, changes in the chromatin accessibility of monocytes were observed 6 days after RIG-I activation, including an enhanced accessibility of transcription factor binding motifs such as STAT1:STAT2, which act downstream of the type-I IFN receptor (IFNAR). Nonetheless, repetitive IFN-β stimulation could not recapitulate the effect of repetitive RIG-I stimulation. Altogether, RIG-I activation induces a persistent “alert-state” at the cellular level, enabling a stronger response upon subsequent immune stimulation, with important consequences for the clinical application of RIG-I agonists in immune therapies.

## RESULTS

### Repetitive 3pRNA stimulation enhances RIG-I signaling at the mRNA and protein levels in human monocytes affording non-specific antiviral protection

To investigate whether RIG-I activation can induce innate immune priming and training, we stimulated monocytes on day 0 and/or day 6 after isolation using a specific RNA RIG-I agonist (3pRNA) previously developed by our group(Schlee *et al*, 2009). Here, we based our experimental approach on other studies on trained immunity(Saeed *et al*, 2014; Bekkering *et al*, 2016b) and compared a single treatment on day 6 (unstimulated followed by RIG-I agonist, UR) to repetitive treatment on days 0 and 6 (RIG-I agonist followed by RIG-I agonist, RR) using monocytes from healthy blood donors (Fig. 1A). In addition to this standard protocol, we characterized potential long-term effects by analyzing cells stimulated on day 0 but not day 6 (RIG-I agonist followed by no stimulation, RU), and we used the transfection of non-stimulatory RNA (UC) on day 6 as a negative control. Moreover, due to the poor cell viability of human monocytes in 6-day culture (data not shown), monocytes were not cultivated in human serum as in previous studies(Saeed *et al*, 2014; Bekkering *et al*, 2016b) but rather in 10% FBS supplemented with 100 pg/mL human M-CSF.

**Figure 1.**
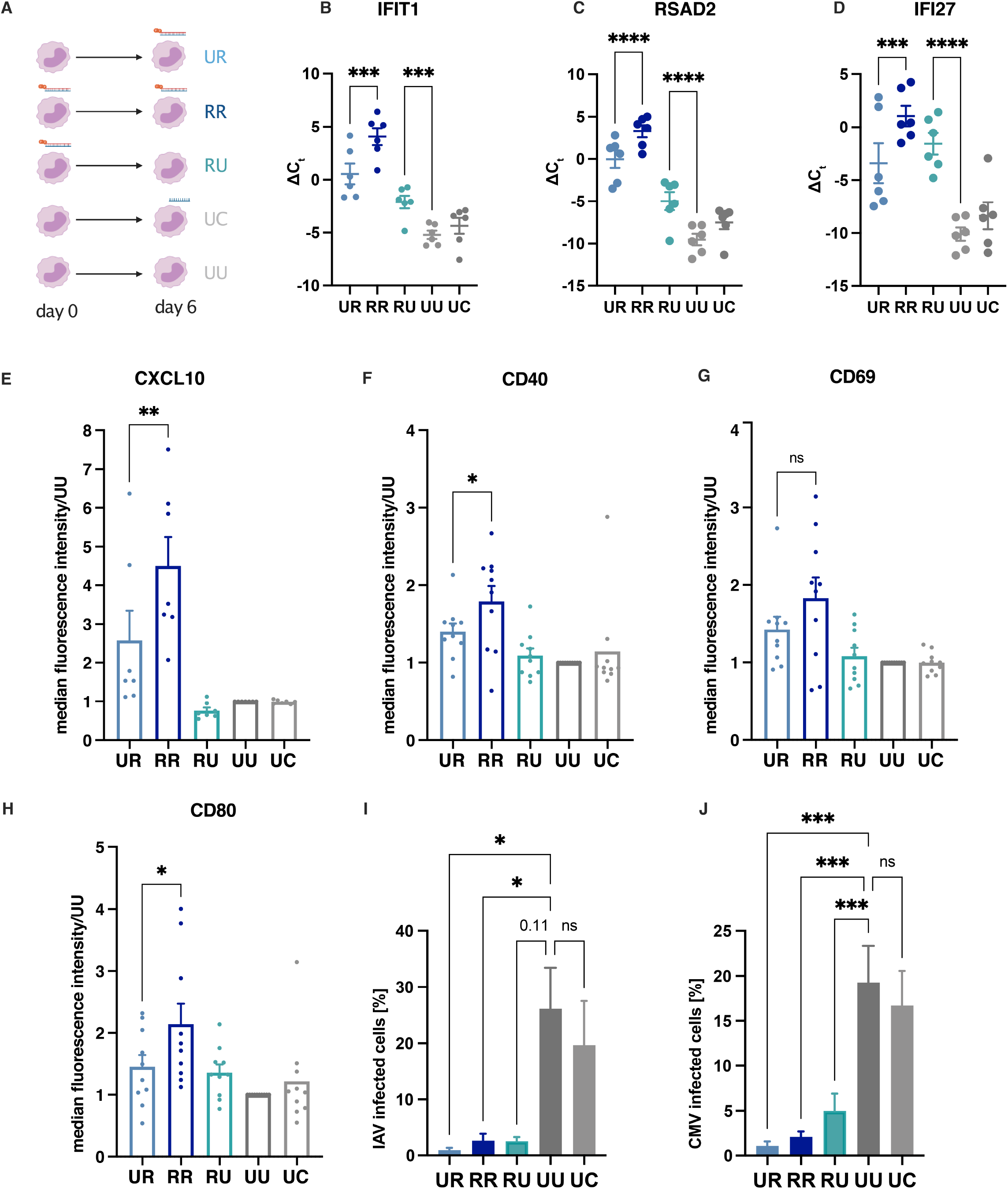
Repetitive 3pRNA stimulation enhances RIG-I signaling at the mRNA and protein levels in human monocytes affording non-specific antiviral protection. (A) Schematic overview of the experimental setting. Monocytes were either unstimulated (U) or stimulated with 3pRNA (R) or control RNA (C) at day 0 and day 6. (B–D) Interferon-stimulated gene expression (*IFIT1, RSAD2, IFI27*) was determined using RT-PCR 6h after stimulation at day 6. Graphs show the mean ± SEM (n=6) for ΔCt (*GAPDH* normalized). (E-H) CXCL10 production (mean ± SEM, n=7) and CD40, CD69, CD80 expression (mean ± SEM, n=10) were measured by FACS 18h after stimulation at day 6. Shown is the median fluorescence intensity relative to UU for individual donors. (I-J) Monocytes were infected with Influenza A virus (IAV, I) and Cytomegalovirus (CMV, J) 24h after d6-stimulation. Infection was quantified by flow cytometry using staining for nuclear protein 8h post infection (p.i., I) and mNeonGreen expression 24h p.i. (J). Graphs are plotted as the mean ± SEM of individual donors (n=4). P-values were determined by one-way ANOVA using Holm-Šídák correction (B-D,J), Wilcoxon matched-pairs signed rank test (E-H) and Friedman’s test using Dunn’s correction (I). ns= not significant, *p ≤ 0.05, **p ≤ 0.01, ***p ≤ 0.001, ****p ≤ 0.0001

As expected, RIG-I activation (UR) robustly induced mRNA transcription of the ISGs, *IFIT1, RSAD2*, and *IFI27* (Fig. 1B-D) and the cytokine CXCL10 (Fig. S1A). Moreover, we could also observe that repetitive (RR) activation of RIG-I induced a stronger ISG response than single (UR) RIG-I activation (Fig. 1B-D) with a similar but not significant tendency for CXCL10 (Fig. S1A). Of note, *IFIT1, RSAD2* and *IFI27* also remained significantly upregulated in RU samples (Fig. 1B-D), 6 days after activation.

Since type-I IFN can limit protein translation and cell viability as part of the antiviral response(Li *et al*, 2015), we also investigated the effect of single and repetitive RIG-I stimulation at the protein level by measuring the antiviral cytokine CXCL10 with ELISA (Fig. S1C). To avoid biases due to differences in cell viability, the CXCL10 levels obtained in ELISA were normalized using cell counts of DAPI-stained nuclei from controls without restimulation (UU,RU) (Fig. S1D, E). Similar to mRNA induction, we found significantly higher amounts of CXCL10 protein release after repetitive RIG-I stimulation when compared to single stimulation (UR), but also six days after single stimulation (RU). Moreover, potential confounding effects resulting from the repetitive administration of the transfection reagent were investigated by comparing CXCL10 levels after single RIG-I stimulation (UR,RU) to two different control RNAs, the single-stranded antisense oligonucleotide of the dsRNA RIG-I ligand (AR,RA) and a non-stimulatory 21-mer CA sequence (CR,RC). The use of these controls demonstrated that transfection itself did not enhance RIG-I signaling (Fig. S1F). Since cell number differences have also been described as a possible mechanism of putative training effects(Garcia-Valtanen *et al*, 2017), we also assessed CXCL10 production at the single-cell level using flow cytometry. Again, we found a significantly higher CXCL10 induction after repetitive stimulation (Fig. 1E). In addition, we examined the cell-surface expression of the activation markers CD40, CD69, CD80, and CD83 (Fig. 1F-H, Fig. S1G,H)(Rusinova *et al*, 2013). The cell-surface expression of all three activation markers was increased, and CD40 and CD80 were significantly enhanced by repetitive RIG-I stimulation (RR vs. UR).

To assess whether single and repetitive RIG-I activation also induce an unspecific protection against reinfection as described for other published trained immunity agents(Netea *et al*, 2016), we infected monocytes with both DNA and RNA viruses, using a H1N1 Influenza A strain (IAV), as a negative-strand (-ss)RNA virus activating RIG-I signaling(Opitz *et al*, 2007), as well as Cytomegalovirus (CMV), a DNA virus which does not typically activate RIG-I(Marques *et al*, 2018). To measure CMV replication, we used a CMV containing a mNeonGreen (mNG) reporter under the control of the HMCV major immediate early promoter (Kasmapour *et al*, 2017). Monocytes were infected with virus 24h after stimulation on day 6(d6). Infection rate was quantified by flow cytometry using intracellular staining for viral nucleoprotein (IAV) and mNG expression for the genetically modified CMV virus (Fig. 1I,J Fig. S1I-K). In the IAV infection model, single and repetitive 3pRNA stimulated monocytes (UR, RR) were significantly less infected when compared to unstimulated cells, whereas single 3pRNA stimulation at day 0 had a marked, but not statistically significant, protective effect (p-value 0.11, Fig. 1I). In the CMV infection model 3pRNA stimulation decreased infection rate after single and repetitive stimulation (RU, RR, UR, Fig. 1J), indicating 3pRNA induced unspecific protection against viral pathogens. Altogether, these results demonstrate that repetitive RIG-I activation increases ISG induction, cytokine release and monocyte activation, with an enhanced unspecific antiviral protection at the cellular level.

### Repetitive 3pRNA stimulation induces an immunological alert state including sensitization of P2RX7 mediated NLRRP3 inflammasome activation

In order to characterize the global transcriptional changes induced by repetitive RIG-I activation, we performed 3’mRNA sequencing with monocytes 24hrs after 3pRNA stimulation at day 0 (untreated: U, 3pRNAstimulation: R) and with day 6 monocytes treated as shown in Fig.1A. Overall, we found broad changes in monocyte RNA expression, especially after repetitive 3pRNA stimulation (Fig. 2A-C). In the principal component analysis (PCA) (Fig. S2A), PC1, which contributed to 40% of the overall variance, had the greatest variance between repetitive stimulation (RR) and untreated samples (UU), with single 3pRNA stimulation on day 0 and on day 6 (UR, RU) clustering in between. PC2 clustered according to the number of days in culture, reflecting the differentiation from monocytes into a more macrophage-like phenotype.

**Figure 2.**
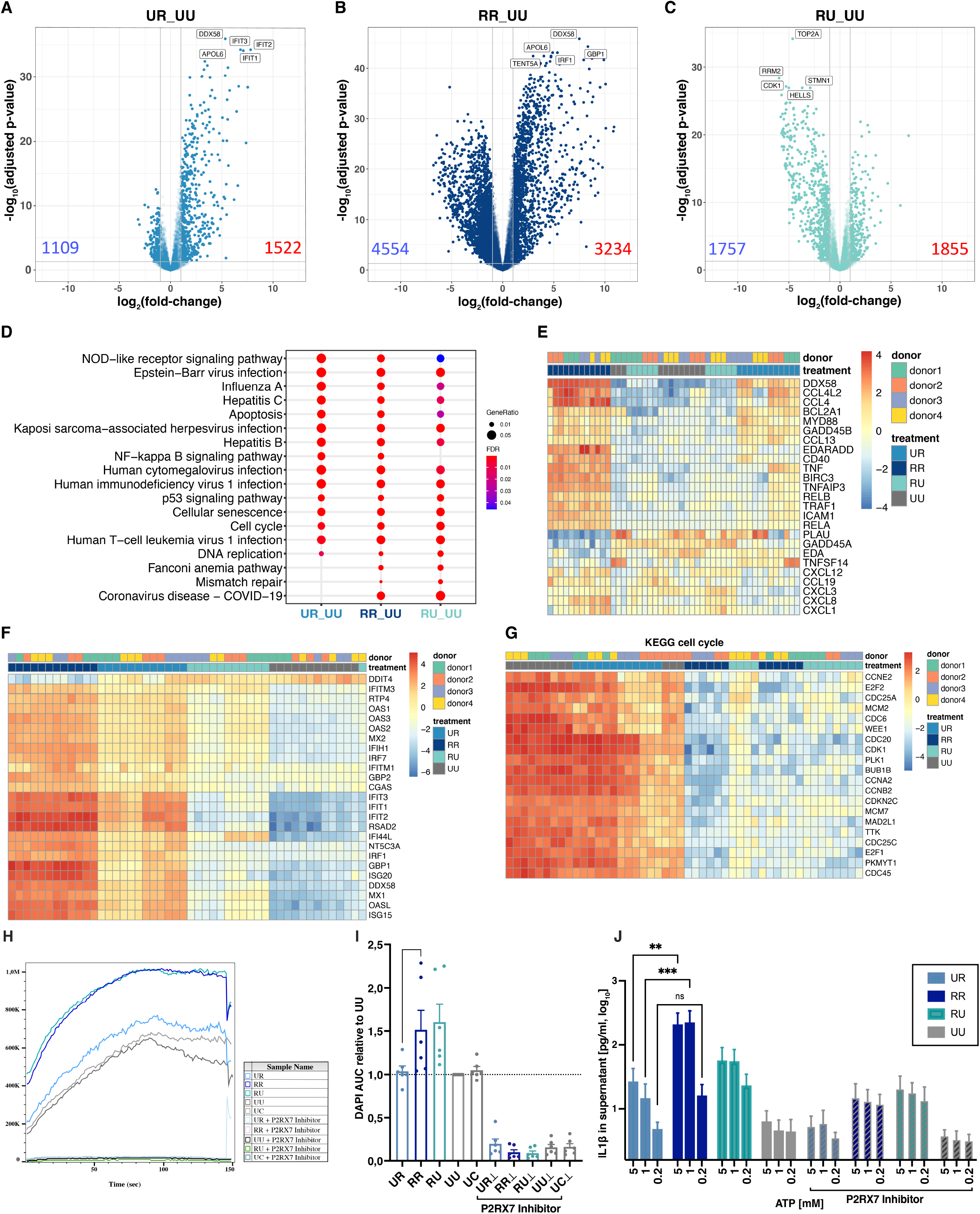
Repetitive 3pRNA stimulation induces an immunological alert state including sensitization of P2RX7 mediated NLRRP3 inflammasome activation. RNA for 3’mRNA sequencing was isolated from monocytes (n=4) harvested 6h after stimulation at day 6. (A-C) Volcano plots with annotation of the top significantly regulated genes are shown for the indicated comparisons. Number of upregulated differentially expressed genes (DEG) shown in red; downregulated DEG shown in blue. (D) Enriched KEGG pathways determined using DEG for the indicated comparisons as the input. (E-G) The most variable genes included in the KEGG-pathway NF-κB signaling (hsa04064, E), in antiviral interferon stimulated genes (from Schoggins & Rice 2011, F) and KEGG-pathway cell cycle (hsa04110, G) are shown in the heatmap. (H-I) P2RX7-mediated DAPI uptake by Bz-ATP stimulation 6h after secondary stimulation, as determined by flow cytometry. (H) Median fluorescence intensity (90% percentile range) is plotted over time for a representative donor. (I) Uptake was quantified as the area under the curve (AUC, 30 sec from Bz-ATP) relative to UU and plotted as the mean ± SEM of individual donors (n=6) with p-values determined by ratio-paired t-test. If indicated, P2RX7 inhibitor (A7400003) was added to samples before Bz-ATP exposure. (J) ATP-dependent IL1β release 5h after d6-stimulation is shown for the indicated extracellular ATP concentrations after normalization for cell numbers. If indicated, P2RX7 inhibitor (A7400003) was added 1h prior to ATP exposure. Graph is plotted as the mean ± SEM of individual donors (n=8) with P-values determined by two-way ANOVA with Šidák correction.

As expected, we found that a substantial number of genes (2631) were differentially expressed (false discovery rate adjusted p-value, abbr. FDR<0.05) after single 3pRNA stimulation (UR versus UU, abbr. as UR_UU), including many type-I and type-II ISGs known to be involved in RIG-I signaling (Fig. 2A, Fig. S2B). However, repetitive stimulation (RR) resulted in a further increase in differentially expressed genes (DEG), in comparison both with unstimulated cells (RR_UU, DEG 7788, Fig. 2B, Fig. S2B) and with cells stimulated on day 0 (RR_RU, DEG 6733, Fig. S2B). While the majority of the DEG from single stimulation on day 6 (UR) were a subset of those differentially expressed after repetitive stimulation (only 2% unique in UR_UU out of total DEG), 39% of DEG were significant in RR_UU but neither in UR_UU nor in RU_UU (Fig. S2C).

To examine whether the differences in gene expression resulted from the induction of unique genes or the stronger induction of a similar response, we plotted the logarithmic fold-changes (LFC) for repetitive and single stimulation (RR_UU vs. UR_UU) and performed Spearman correlation analysis and linear regression modeling (Fig. S2D), which revealed a close correlation (r=0.76) and linear slope of 1.9, thus demonstrating that repetitive stimulation induces a quantitatively stronger but qualitatively similar transcriptional response compared to single stimulation on day 6. Euclidean cluster distancing of the 20 most significantly regulated genes from RR_UR resulted in treatment-dependent clustering (Fig. S2E), with a clear induction of type-I or type-II ISGs (Interferome). Moreover, KEGG pathway analysis revealed, as expected, the enrichment of several pathways associated with viral infection, including Influenza A, Hepatitis B and C, and COVID-19 (Fig. 2D), as well as changes in several proliferation- and survival-associated pathways, including the induction of p53 signaling, cellular senescence, apoptosis, and modulation of the cell cycle.

Whereas most viral pathways were enriched in all three 3pRNA stimulated conditions (UR, RR, RU), the NFκB pathway was not enriched for RU_UU. Euclidean cluster distancing of genes included in the NFκB pathway led to clustering of RU with UU, with a strong induction in RR when compared to UR indicating immune training rather than a priming effect for 3pRNA induced proinflammatory genes (Fig. 2E). In contrast, most antiviral ISGs(Schoggins & Rice, 2011) were enhanced in RU, indicating immune priming (Fig 2F). Accordingly, GO-term enrichment analysis found the terms *response to viruses* and *type I interferon signaling* to be significantly enriched for single (UR, RU) and repetitive stimulation (Fig. S2F). In total, we found 1854 genes with training effect (significantly upregulated in RR_UU, LFC RR_UU>UR_UU, not DE in RU_UU) and 1168 with a priming effect (significantly upregulated in RR_UU, LFC RR_UU>UR_UU, significantly upregulated in RU_UU). However, only 79 DEG were tolerized (significantly upregulated in UR_UU, LFC RR_UU<UR_UU) by repetitive stimulation without an enriched KEGG-pathway (Fig S2G).

Functional overrepresentation analysis was subsequently performed with the four main clusters identified by expression similarity analysis. Here, we found several of the previously described viral infection pathways (Fig. S2H) in Cluster A, which included genes which were upregulated in day 6 3pRNA (UR, RR) stimulated samples. Genes in cluster B and C were downregulated upon day 6 stimulation, most likely reflecting interferon mediated inhibition of transcription. Genes associated with the cell cycle were overrepresented in Cluster D, which primarily consisted of genes downregulated by training (RR, RU). This cell cycle regulation could be confirmed by Euclidean cluster distancing of genes included in the cell-cycle pathway (hsa04110), which led to clustering dependent on the status of 3pRNA stimulation at day 0 (Fig. 2G, RR, RU). This is in line with the relative reduction in cell number observed for 3pRNA-trained cells (Fig. S1D) and with previous studies on IFN-induced cell-cycle arrest(Sangfelt, 2000).

NOD-like receptor signaling was among the most significantly enriched KEGG pathways after repetitive 3pRNA treatment in monocytes (Fig. 2D). When we examined the genes linked to this pathway in more detail (Fig S2I), we found that 3pRNA stimulation significantly upregulated the expression of P2X purinoceptor 7 (*P2RX7*), an ATP-gated ion channel that induces potassium efflux and can thus activate the NLRP3 inflammasome (Surprenant *et al*, 1996; Solle *et al*, 2000; Pelegrin, 2021). Since IFN is generally regarded as a negative regulator of NLRP3 activation(Guarda *et al*, 2011b; Reboldi *et al*, 2014; Dang *et al*, 2017), we were intrigued by this finding. RT-PCR of *P2RX7* and key inflammasome components after IFN*β*, LPS, and 3pRNA stimulation (Fig. S2J-M), confirmed that *P2RX7* is significantly induced by 3pRNA, TNF and LPS but not IFN*β*. In line with the mRNA sequencing results (Fig. S2N-Q), 3pRNA stimulation at day 6 did not induce *NLRP3* or *IL1B* expression. To assess the functional relevance of these observations, we measured P2RX7 activation-dependent intracellular DAPI uptake via flow cytometry(Proietti *et al*, 2014) after monocytes were stimulated with 3pRNA, as shown in Fig. 1A, and subsequently treated with Bz-ATP, a P2RX7 specific agonist(Surprenant *et al*, 1996; Pelegrin, 2021). DAPI uptake was then quantified by assessing the area under the curve (AUC) relative to untreated samples for each donor. Repetitive 3pRNA stimulation induced faster DAPI uptake than untreated, mock-treated, or single-stimulated cells (Fig. 2H,I, S2R). Surprisingly, RU treatment also accelerated DAPI uptake, although we did not find a significant induction of *P2RX7* at the mRNA level with RU treatment (Fig. S2P). Of note, DAPI uptake was abrogated for all samples in the presence of the P2RX7 receptor antagonist A7400003 (Fig. 2H,I).

Activation of P2RX7 is triggered by high extracellular ATP concentrations and results in NLRP3-inflammasome-dependent IL-1β release from NF-κB-primed macrophages(Pelegrin & Surprenant, 2006; Pétrilli *et al*, 2007), most commonly using activators of TLR2 or TLR4 (Bauernfeind *et al*, 2009). Using cells on day 6 of M-CSF treatment, as in the UU control, we could reproduce priming dependent IL-1β release by showing that ATP induced IL1β release in monocytes primed with the TLR2 agonist Pam3Cys (UP) but not in naive monocytes (Fig. S2S). However, RIG-I activation (UR, RU, and RR), without Pam3Cys treatment, was also sufficient to prime P2RX7-mediated NLRP3 activation in monocytes (Fig. 2J), with RR-treated samples showing the most robust IL1β-response to stimulation with both 5 mM and 1 mM ATP. IL-1β release was decreased for all samples in the presence of A7400003, showing that IL-1β release was P2RX7-dependent, and, as expected, A7400003 did not affect IL-1β release by nigericin-induced potassium efflux (Fig. S2T).

Since these data demonstrate that 3pRNA training does not only enhance subsequent 3pRNA sensing but also sensitizes other host defense mechanisms such as the inflammasome, we then broadly assessed whether TLRs and other pattern recognition receptors (PRRs) involved in viral recognition were differentially expressed upon 3pRNA training (Fig. S2U). Most of these genes were differentially expressed upon repetitive 3pRNA stimulation, and only *TLR5, TLR6* and *MRC1* were downregulated. For trained cells without restimulation (RU), we found 9 DEG, all of which were upregulated, and those with the strongest induction were *MX1, DDX58* (RIG-I), *AIM2*, and *IFIH1* (MDA5). Altogether, these data indicate that 3pRNA stimulation at day 0 does not only alter subsequent RIG-I signaling but also induces multiple antiviral mechanisms resulting in a trained immunologically protective state. Moreover, the lack of immune tolerance induced (Fig. S2G) by RIG-I activation renders it amenable to multiple applications without losing its antiviral capacity at the cellular level.

### 3pRNA stimulation in monocytes induces changes in chromatin accessibility

To assess whether training-induced transcriptional changes could be explained by changes in chromatin accessibility, an assay for transposase-accessible chromatin (ATAC) sequencing was performed in human monocytes at 24h after the first stimulation on day 0 and prior to the second stimulation on day 6 (Fig. S3A). A peak-calling algorithm and sliding-window approach were used for the detection of differentially accessible chromatin regions (DAR)(Feng *et al*, 2020). For both methods, the PCA analysis showed treatment-dependent clustering on day 6 (Fig. S3B, C). We focused on the differentially accessible regions at day 6 (RU_ UU), since this timepoint is the most relevant for repetitive versus single stimulation. The peak-calling approach identified 173 differentially accessible regions (DAR, FDR-adjusted p-values <0.05). Among them, 140 DAR could be assigned to a gene (Fig. S3D,E). 689 DAR were found by the sliding-window approach (Fig. S3D, Fig. 3A), which included 233 DAR related to a known gene, including *CCL2* and *IFI27* (Fig. 3A and B, Fig S3F). Some DAR, such as the ISGs *MX1* and *IFI27*(Schoggins & Rice, 2011), were also upregulated at the mRNA level, whereas others, like the NFκB target gene *CCL2*(Hoesel & Schmid, 2013), were not induced in RU_UU, but rather after d6-stimulation (RR_RU, Fig. S3G-I), revealing that DAR regulation was relevant for both primed and trained genes. Thus, we compared the overlap of DEG at the mRNA level with DAR for each approach and found a relatively small overlap with 126 genes with changes in both transcriptome and chromatin accessibility when comparing RR to UR and 77 common genes using RU_UU (Fig. S3J,K).

**Figure 3.**
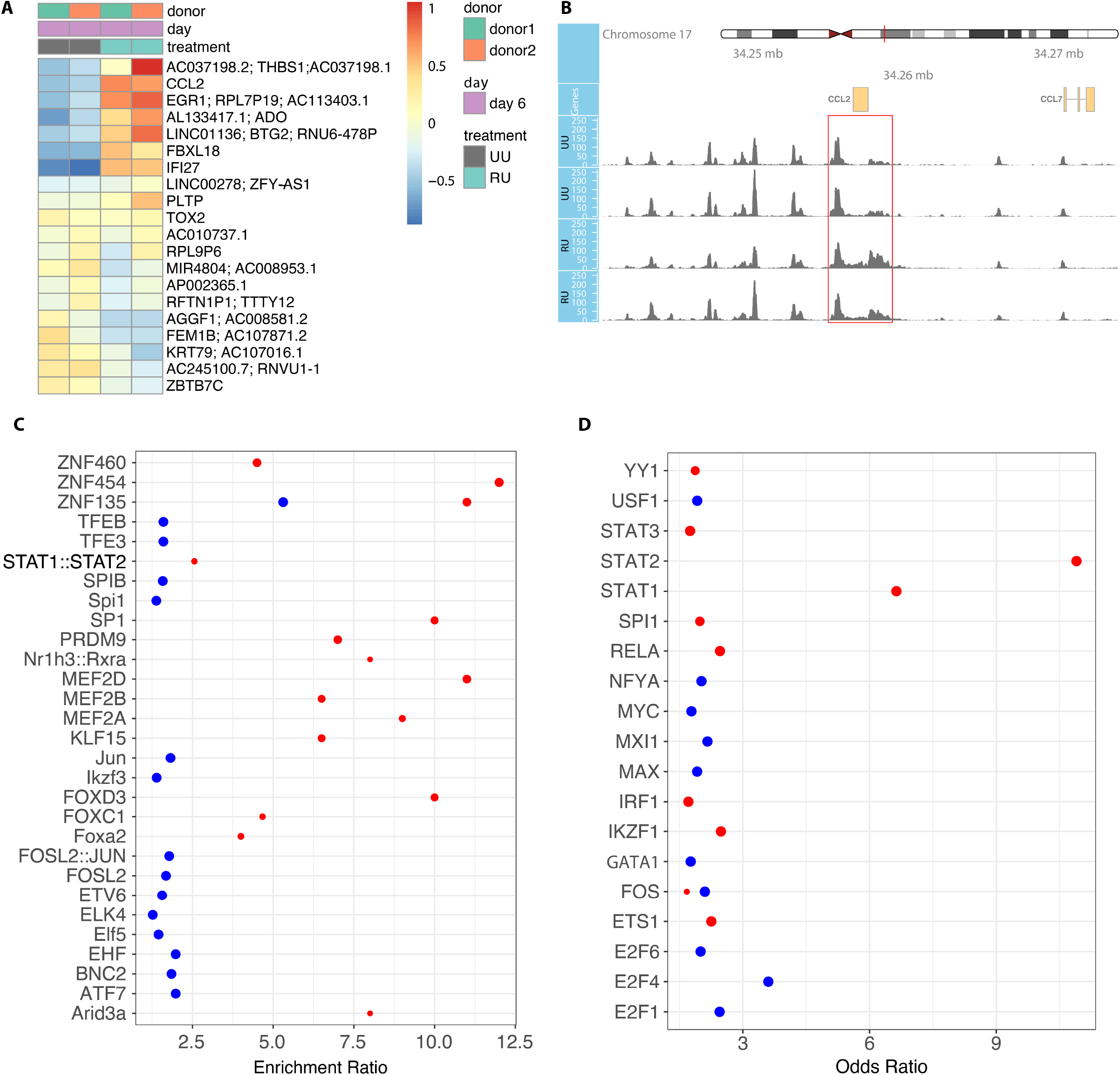
3pRNA stimulation at day 0 induces changes in chromatin accessibility at day 6. Day 0 3pRNA-stimulated (R) and unstimulated (U) monocytes were harvested at day 1 and at day 6 for ATAC sequencing (n=2). (A) Clustered heatmaps show the most significant, gene-annotated, differentially accessible regions (DAR) obtained by sliding-window approach on d6. (B) Reads for individual samples at d6 are shown for *CCL2*, DAR determined by sliding-window approach is highlighted in red. (C) The most enriched transcription factor motifs obtained by SEA using up-(logarithmic fold-change, LFC>0, red dots) and downregulated (LFC<0, blue dots) DAR (sliding window approach) as input are shown. Point size anticorrelates with P-Value. (D) Most significant ChEA3 results for overrepresented transcription factors using DEG in RR vs. UR (mRNA sequencing) as input are shown. Motifs obtained by downregulated DEG (LFC<0) are shown in blue and upregulated DEG (LFC>0) in red with greater point size representing smaller p-value.

Transcriptional differences can also be influenced by changes in the accessibility of transcription factor (TF) binding areas(Klemm *et al*, 2019). Thus, we performed transcription factor motif enrichment analysis of the ATAC sequencing data using DAR for RU_UU obtained by the sliding-window approach (Fig. 3C). Enriched transcription factors motifs (TFM) found using the more accessible regions upon 3pRNA stimulation as input are shown in red, whereas TFM found for less accessible regions are shown in blue. We found several components of the AP-1 group, including FOSL2 and JUN among the top downregulated motifs, indicating that RIG-I training has a differential effect on downstream signaling pathways. In addition, we found several TF motifs of the zinc-finger protein family (ZNF460, ZNF454, ZNF135) as well as STAT1:STAT2 (an essential signal transducer and transcription factor complex downstream of type I, II, and III IFNs(Mogensen, 2019)) to be highly enriched in the more accessible regions (Sup. table 1). Expression of the TFs obtained by the motif enrichment analysis was cross-referenced with the mRNA sequencing data. Here, we found STAT1 and STAT2 to be expressed and upregulated upon repetitive stimulation, whereas the zinc-finger proteins were not expressed under any of the conditions (Fig. S3L). We subsequently used ChEA3, a tool which predicts potential regulatory TFs from a list of genes and used the DEG from the RR_UR comparison to predict key TFs that could be driving the differences between single and repetitive stimulation signaling (Fig 3D, Table S2). Again, we found STAT1 and STAT2 to be highly enriched. We could also detect several TF motifs including for SPI1, SP1, and MEF2A, which were also found in the ATAC motif enrichment analysis (Fig. 3C). Altogether, although we did not generally detect a direct connection between the DAR in close proximity to a gene and corresponding transcriptomic changes at the single-gene level, RIG-I activation induced pronounced changes in chromatin accessibility, which may influence the transcriptome through DAR at regulatory elements.

### Repetitive 3pRNA stimulation enhances and accelerates RIG-I signaling in non-immune cells

As RIG-I is ubiquitously expressed in nucleated cells(Bartok & Hartmann, 2020), we also analyzed the effects of repetitive RIG-I activation in primary human epithelial cells and fibroblasts. These cell types are not only essential immune sentinels at barrier sites(Larsen, 2020) but also have a longer lifespan than monocytes(Sender & Milo, 2021), rendering them particularly subject to repetitive pathogen and, therefore, RIG-I agonist exposure. Using the same experimental approach as in Figure 1 (overview in Fig. 1A), both epithelial cells (Fig. 4A) and fibroblasts (Fig. 4B) showed an enhanced RIG-I response after repetitive stimulation at the mRNA level for the ISGs *IFIT1, RSAD2* and *IFI27*. Moreover, long-term induction of these ISGs could also be observed in epithelial cells and fibroblasts (RU, Fig. 4A,B). 3’mRNA sequencing data indicated a quantitatively stronger RIG-I response after repetitive treatment (RR) compared to single stimulation (UR) in both cell types, with a markedly higher number of DEG in RR_UU (Fig. S4A-D).

**Figure 4.**
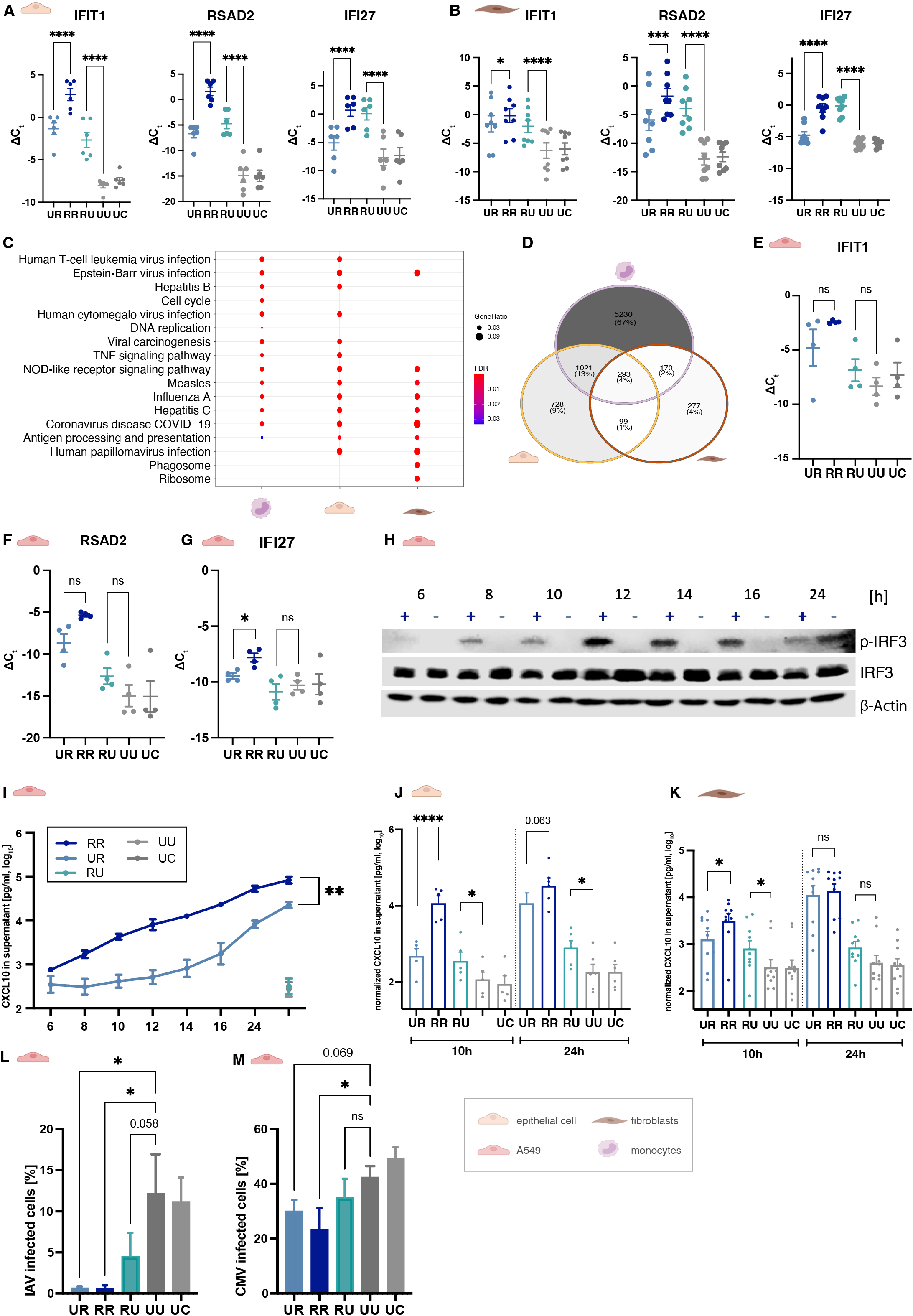
Repetitive 3pRNA stimulation enhances and accelerates RIG-I signaling in non-immune cells. Interferon-stimulated gene expression (ISG, *IFIT1, RSAD2, IFI27*) in epithelial cells (A) and fibroblasts (B) was determined by RT-PCR 6h after d6-stimulation. Graph shows the mean ± SEM (A n=6, B n=8) of ΔCt (*GAPDH* normalized). RNA for 3’mRNA sequencing was isolated 6h after stimulation of epithelial cells (n=3) and fibroblasts (n=4). (C) The top enriched KEGG Pathways using DEG of RR_UR as the input, are shown for monocytes, epithelial cells, and fibroblasts. (D) The overlap of differentially expressed genes (DEG) for RR_UR is shown for monocytes, epithelial cells, and fibroblasts. (E-G) ISG expression (*IFIT1, RSAD2, IFI27*) in A549 was determined by RT-PCR 6h after d13-stimulation. Graph shows the mean ± SEM (n=4) of ΔCt (*GAPDH* normalized). (H) IRF3 phosphorylation (p-IRF3) after 3pRNA treatment of trained (+) and untrained (-) A549 cells was quantified over time by western blot. (I) Supernatants of 3pRNA-treated A549 cells were collected after the indicated time points. CXCL10 was quantified by ELISA and plotted as the mean ± SEM of independent experiments (n=3). (J,K) Supernatants of epithelial cells (J) and fibroblasts (K) were collected 10h and 24h after stimulation at day 6. CXCL10 was quantified by ELISA, normalized for cell number and plotted as the mean ± SEM of individual donors (J n=5, K n=10). (Z, AA) A549 cells were infected with Influenza A virus (IAV, Z) or Cytomegalovirus (CMV, AA) 24h after d6-stimulation. Infection was quantified by flow cytometry 8h (IAV) and 24h (CMV) post infection. Graphs show mean ± SEM of individual donors (Z n=3, AA n=4). P-values were determined by Friedman test using Dunn’s correction (L,M), one-way ANOVA (A,B,E,F,G,J,K) and two-way ANOVA (I) using Holm-Šídák correction. ns = not significant; *p ≤ 0.05, **p≤0.01, ***p ≤0.001, ****p ≤0.0001.

As observed for monocytes (Fig. S2D), epithelial cells and fibroblasts demonstrated the highest number of DEG between RR and UU in the comparison to UR_UU and RU_UU (Fig. S4E,F). However, the overall number of DEG were lower in both cell types (RR_UU: 2851 fibroblasts, 2184 epithelial cells; compared to 7788 in monocytes). Accordingly, less than 77 epithelial and 527 fibroblast genes were differentially regulated in the comparison UR_UU, and, of these, only 8 epithelial and 45 fibroblast genes were uniquely induced by single stimulation (Fig. S4E,F). Although KEGG pathway analyses for the main comparisons (UR_UU, RR_UU and RU_UU) found an enrichment of similar pathways compared to the monocyte data (Fig. 4C, Fig. 2D, Fig. S4G,H), only 4% of the DEG when comparing RR to UR were differentially expressed in all three cell types, and 67% were uniquely enriched in the monocyte data (Fig. 4D). To exclude that the rather small overlap of common inducible genes was due to cell-type specific transcriptomes, we analyzed the overlap of all expressed genes in the dataset, resulting in 66% of genes being commonly expressed (Fig. S3I). Among the unique DEG in epithelial cells, we found that the type III IFN genes *IFNL2* and *IFNL3* were significantly upregulated. Moreover, functional enrichment analysis of the DEG showed an overrepresentation of genes associated with the cell cycle for the monocyte data set but not for the other cell types (Fig. 4C, fourth lane). In contrast to what was observed in monocytes (Fig 2G), heatmaps plotted with cell-cycle genes did not cluster in a treatment-dependent manner for fibroblast or epithelial-cell data sets (Fig. S4J, K), and, in line with these observations, RIG-I activation in fibroblasts and epithelial cells did not substantially affect cell number (Fig. S4L, M). Thus, a substantial difference in DEG between monocytes and fibroblasts and epithelial cells is the lack of cell cycle regulation downstream of RIG-I activation in these cells, in contrast to RIG-I stimulated monocytes. Unlike what was observed in monocytes (Fig. 2E), NFκB signaling was only slightly induced by single 3pRNA stimulation. A clear upregulation of this pathway was only shown after repetitive 3pRNA stimulation in epithelial cells, indicating that training is critically required for significant NFκB induction in these cells (Fig. S4N). In contrast, fibroblasts did not demonstrate significant NFκB induction even after repetitive stimulation (Fig. S4O).

To determine the duration of altered signaling, as well a potential preservation of TI effects across cell division, we used A549 cells, an alveolar basal epithelial cell line derived from a human adenocarcinoma(Giard *et al*, 1973). A549 cells demonstrated enhanced ISG expression after repetitive stimulation at the mRNA level (Fig. S4P-R) that resembled the response seen with the primary epithelial cells. Using 13 and 28 days instead of the previously described d6 interval we found a slight increase in ISG signaling but significant only for *IFI27* after repetitive stimulation after 13 days (Fig. 4E-G), and no differences in-between single and repetitive stimulation after 28 days (Fig S4S-U), demonstrating the temporary nature of TI in this cell type.

We then investigated whether RIG-I training affected the speed of RIG-I signaling. A time course of IRF3 phosphorylation after d6-3pRNA stimulation showed a rapid IRF3 response in 3pRNA-trained cells (RR cells) with the maximum phosphorylation around 12–14h, whereas untrained cells (UR cells) demonstrated maximum IRF3 phosphorylation around 24h after stimulation (Fig. 4H, controls in Fig. S4V). Similarly, repetitively stimulated cells showed a more rapid CXCL10 release, as well as a higher cumulative CXCL10 level compared to single 3pRNA stimulation (Fig. 4I). To confirm these findings in primary cells, we examined CXCL10 levels 10h and 24h after RIG-I stimulation for epithelial cells and fibroblasts (Fig. 4J,K). We found significantly higher CXCL10 levels in epithelial cells and fibroblasts 10h after repetitive RIG-I stimulation (RR) compared to previously unstimulated cells (UR). However, no significant difference was seen when CXCL10 levels were determined in the supernatants at 24h. These results demonstrate a faster rather than a stronger response in trained epithelial cells and fibroblasts. We then investigated whether monocytes demonstrated a similar difference in RIG-I activation kinetics. Here, CXCL10 levels after restimulation were also significantly enhanced both at 10h and 24h demonstrating a stronger response at both early and late time points after stimulation (Fig. S4W, Fig. S1C). In line with the upregulation of PRR upon 3pRNA stimulation in monocytes (Fig S2U), repetitive stimulation of epithelial cells and fibroblasts also strongly induced *DDX58* (RIG-I), *IFIH1* (MDA5), *MX1*, and *TLR3* (Fig. S4X,Y), albeit a lower number of expressed PRR. We then examined the effect of RIG-I stimulation in A549 in the established IAV and CMV infection models (Fig. 4L,M Fig. S4Z). As in monocytes, prior RIG-I stimulation reduced the percent of infected cells for both viruses, although this did not reach statistical significance for stimulation on day 0. Here, repetitive RIG-I stimulation enhanced antiviral protection and led to the clearest reduction in IAV and CMV infected cells, providing a further indication that repetitive RIG-I stimulation trained rather tolerizing the antiviral response.

### Enhanced RIG-I signaling is STAT1 dependent, yet type-I IFN stimulation alone is not sufficient for training

Due to the enhanced motif accessibility of STAT1 and STAT2 in the ATAC sequencing data (Fig. 3C) and the previously published role of STAT1 in *β*-glucan training (Ifrim *et al*, 2015), we decided to investigate the contribution of STAT1 signaling to RIG-I-induced immune training using *STAT1*^-/-^ A549 cells (Fig. S5A). As in previous experiments, the cells on d0 and d6 were either unstimulated (U) or stimulated with 3pRNA (R) or control RNA (C). Similar to the data obtained with primary cells, RR-treated wildtype A549 cells showed a significantly higher response compared to UR treated samples for most of the analyzed genes (Fig 5A-D). However, this effect was abrogated in *STAT1^-/-^* A549 cells, not only for ISGs such as *IFIT1* but also for the NF-κB-inducible gene *IL6*(Hoesel & Schmid, 2013) (Fig. 5A-D).

**Figure 5.**
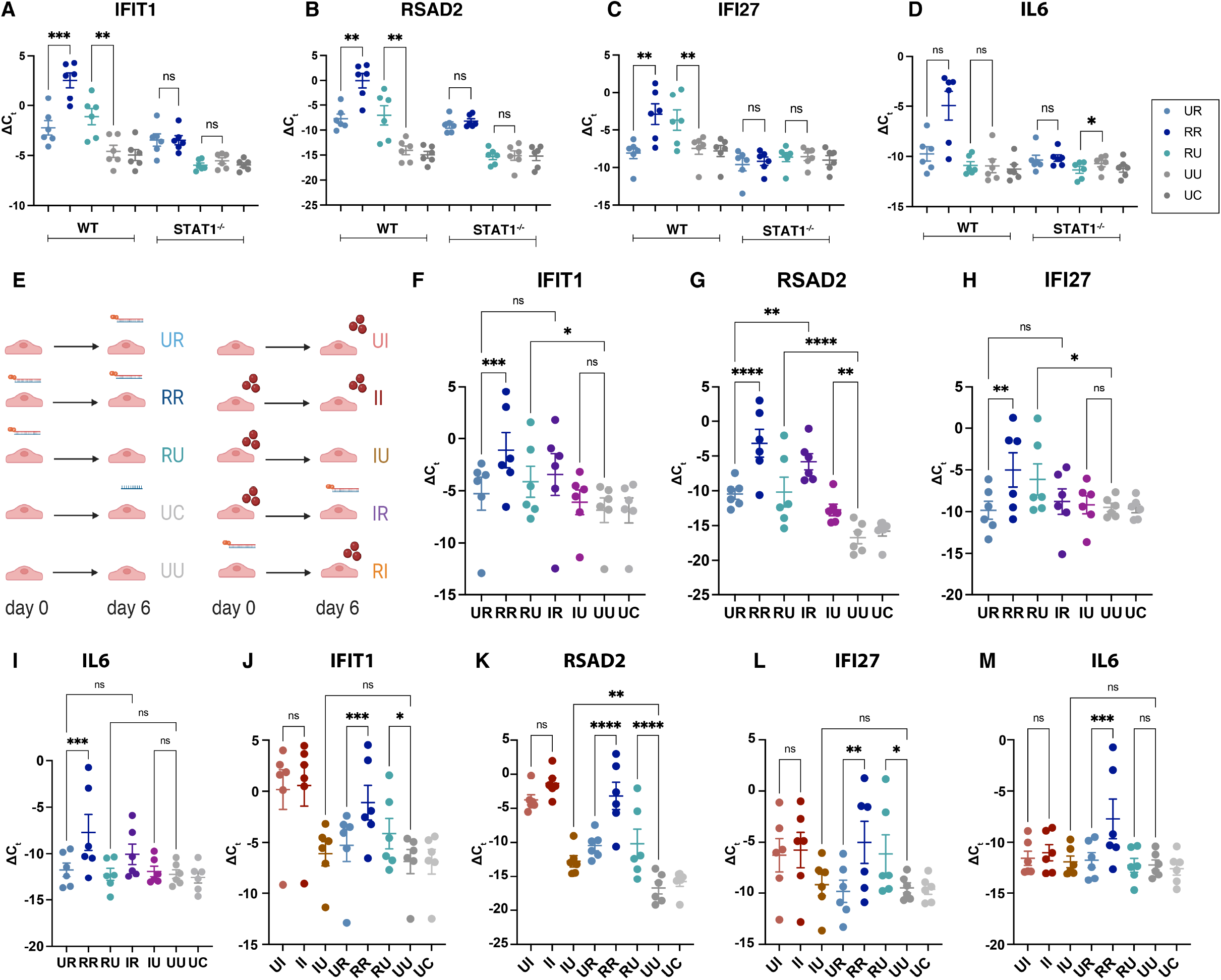
Enhanced RIG-I signaling is STAT1 dependent but interferon β stimulation alone is not sufficient for training. (A-D) RIG-I signaling was determined using RT-PCR 6h after indicated stimulations of wildtype (WT) and STAT1^-/-^ A549 cells. Graphs show the mean ± SEM of -ΔCt (*GAPDH* normalized, n=6) for indicated genes. (E-M) RIG-I signaling was determined by RT-PCR 6h after A549 WT were either unstimulated (U) or stimulated with 3pRNA (R), Interferon β (IFNβ= I, 100,000 U/ml) or control RNA (C) at day 0 and day 6. (E) Schematic overview of the experimental setting. (F-M) Graphs show the mean ± SEM of -ΔCt (*GAPDH* normalized, n=6) for indicated genes. (F-I) compare 3pRNA and IFNβ training, whereas repetitive IFNβ stimulation is shown in (J-M). P-values were determined by one-way ANOVA using Holm-Šídák correction. ns = not significant, *p ≤ 0.05, **p ≤ 0.01, ***p ≤ 0.001, ****p ≤ 0.0001

Since this demonstrated that IFN signaling was necessary for RIG-I TI, we then investigated whether type-I IFN treatment alone would be sufficient to induce the observed training and priming effects (Fig. 5E-I). For these experiments, we initially stimulated cells with recombinant IFNβ at a concentration similar to the highest amount released by 3pRNA stimulation (100,000 U/ml, Fig. S5B) instead of using the RIG-I agonist. For this amount of IFNβ, we could observe priming (IU_UU) and training (IR_UR) for RSAD2 only (Fig. 5G). However, in comparison with RIG-I, the overall effect of IFNβ training and priming was less pronounced. Moreover, in contrast to prior RIG-I activation, no effect of IFNβ on innate immune memory could be observed for IFIT1, IFI27 or IL6 (Fig. 5F,H,I), indicating that RIG-I activation affects a broader number of genes than IFNβ alone. Similar results were found when comparing repetitive IFNβ stimulation (II_UI) to repetitive RIG-I stimulation directly(Fig 5J-M). Despite the observed long-term induction of RSAD2 (IU_UU) we did not find any significant difference between single and repetitive IFNβ stimulation (Fig 5J-M).

Conversely, we then investigated whether IFNβ stimulation on day 6 could be enhanced by prior 3pRNA stimulation (RI) and compared this to repetitive stimulation with IFNβ (II). Here, prior RIG-I training also resulted in a slight increase in ISG induction compared to IFNβ treatment alone (RI_UI, Fig. S5C-F, RI vs. II fold change RSAD2 1.2, IFIT1 1.7), which was again only significant for RSAD2 (Fig. S5D). To rule out a potential ceiling effect for IFNβ activity, we then primed cells with a lower concentration of IFNβ (2000 U/ml), which gave comparable results (Fig. S5G-J). Altogether, our data indicate that, while type-I IFN signaling via its key signaling molecule STAT1 is necessary for RIG-I mediated training, type-I IFN stimulation alone is not sufficient to compensate for direct RIG-I activation.

## DISCUSSION

In the present work, we demonstrate, for the first time, that RIG-I activation can induce trained immunity (TI). Immune priming and training responses were observed in monocytes, epithelial cells and fibroblasts at the protein and RNA levels, including both anti-viral genes and pathogen recognition receptors. In line with these findings, 3pRNA-mediated priming and training afforded effective, non-specific antiviral protection against CMV and IAV infection. Changes in chromatin accessibility were confirmed in ATAC sequencing, which indicated a role for STAT1:STAT2 accessibility in RIG-I mediated TI. However, although STAT1 signaling was indeed required for RIG-I induced TI, interferon-β mediated STAT1 activation did not recapitulate the effect of RIG-I TI. Furthermore, we show, for the first time, that accelerated kinetics of signal transduction is a component of the trained immunity phenotype. Taken together, our data demonstrates that RIG-I activation induces an immunologically long-term “alert state”, licensing heightened immune reactivity and resistance to infection, reminiscent of what has been described for *β*-glucan and BCG training.

Numerous publications have reported non-specific immune protection after administration of nucleic acid agonists *in vivo*, including activators of RIG-I, TLR3, TLR7, TLR9. As with BCG(Starr *et al*, 1976; Garly *et al*, 2003; Kleinnijenhuis *et al*, 2012), these studies often focus on the protective effects in very young or immune-compromised hosts which exhibit limited adaptive immune responses (Verthelyi *et al*, 2003; Ito *et al*, 2005; Pedras-Vasconcelos *et al*, 2006; Ribes *et al*, 2014) and are thus strongly suggestive of innate immune memory. Nonetheless, only one study to date has specifically investigated the immune training and tolerance induced by nucleic-acid (NA) immune sensors. Using the standard monocyte model(Bekkering *et al*, 2016a), this study investigated endosomal (TLR3,7-9) but not cytosolic NA receptors and reported dose-dependent induction of training or tolerance for TLR3 and TLR7/8, using poly(I:C) and R848, while the TLR9 agonist CpG had no memory effect. In contrast, in our study, only a small subset of genes was tolerized upon RIG-I activation with 3pRNA (Fig S2G), and these tolerized genes were not directly related to innate immune signaling or effector functions.

Although this may seem to indicate a fundamental difference between endosomal and cytosolic sensing of nucleic acids, these results should be interpreted with caution due to the cellular model employed. TLR9 is not expressed in human monocytes (Fig. S2U and (Katona & Lindskog, 2022)), which provides a clear explanation for why no training effects were observed for the TLR9 ligand CpG. Likewise, TLR3 and TLR7 are only expressed at low levels in resting monocytes, although we found that both TLR3 and TLR7 are upregulated by RIG-I training (Fig. S2U). Thus, in monocytes, it seems likely that poly(I:C) would, at least initially, activate RIG-I, MDA5, PKR, and OAS, as well as potentially ZBP1, DHX and DDX proteins(Bartok & Hartmann, 2020), and, indeed, the interaction of these receptors and their respective activation thresholds may explain the dose-dependent effects observed. Likewise, R848 is an agonist of both TLR7 and TLR8. In monocytes, its activation does not induce IFNα but rather a pro-inflammatory cytokine profile including IL12p70 and only low levels of IFNβ(Ablasser *et al*, 2009; Bergstrøm *et al*, 2015). Of note, these effects are in stark contrast to the *in vivo* application of these ligands in mice and humans, which activate TLR7 and TLR9 in pDCs, inducing high levels of IFNα, as well as TLR7 and TLR9 on macrophages and DCs in mice(Hartmann, 2017; Bartok & Hartmann, 2020), inducing the additional release of pro-inflammatory cytokines. Similarly, poly(I:C) can also activate TLR3 on fibroblasts and DCs *in vivo* in addition to many other dsRNA receptors(Muzio *et al*, 2000; Bartok & Hartmann, 2020). Thus, creating appropriate *in vitro* cellular models for NA-sensing TLR TI *in vivo* is exceptionally challenging.

In contrast, RIG-I is virtually ubiquitously expressed in mammalian cells; both *in vivo* and *in vitro* studies have demonstrated antiviral protection after 3pRNA administration(Liu *et al*, 2016; Coch *et al*, 2017a; Mao *et al*, 2021; Marx *et al*, 2022; Schwab *et al*, 2022b, 2022a), and specific ligands are available(Junt & Barchet, 2015; Feld & Schuberth-Wagner, 2022), all of which render RIG-I activation particularly amenable to *in vitro* TI models, including both the standard monocyte model and non-immune cells. However, to our knowledge, no study to date has specifically investigated RIG-I-induced innate immune priming or training. Thus, our current study provides not only new insight into RIG-I signaling but also key information to understand what was previously observed in infection models after RIG-I ligand application. Our data demonstrate that many antiviral effectors are primed by RIG-I (Fig. 2C, D), remaining upregulated after 6 days, and thus providing a direct, yet non-specific, block of viral entry and replication, which likely contributes to the effectiveness of RIG-I stimulation against a broad variety of both RNA and DNA viruses. Moreover, we observed that innate immune training enhanced both the strength and the speed of the antiviral response upon RIG-I restimulation. RIG-I training led to more rapid IRF3 phosphorylation and CXCL10 release after the second stimulation (Fig. 4G,I). Accordingly, enhanced chromatin accessibility was not only found for STAT1:STAT2 transcription factor motif binding sites but also around transcription start sites of antiviral genes such as MX1 (Fig S3E), an antiviral effector protein which we have previously shown to be essential in RIG-I mediated defense against IAV(Schwab *et al*, 2022c). As several recent studies on SARS-CoV-2 have demonstrated(Cheemarla *et al*, 2021; Decker, 2021; Lopez *et al*, 2021), the speed of the innate immune response is critical to an effective host defense, particularly since SARS-CoV-2 and many other viruses code for proteins that delay ISG induction(Shemesh, 2021; Fung *et al*, 2022). Given that SARS-CoV-2 and IAV both activate RIG-I like receptors(Rehwinkel *et al*, 2010), these findings may also explain why RIG-I is a particularly effective prophylaxis against these two viruses(Marx *et al*, 2022; Mao *et al*, 2021b; Coch *et al*, 2017b).

Our study was designed to distinguish between the priming and training effects of RIG-I, as advocated by Divangahi et. al(Divangahi *et al*, 2021b). Analyzing RNA transcription at day 6 after stimulation(Bekkering *et al*, 2016a), we found that 3pRNA-treated cells (RU) still actively transcribed certain ISGs such as IFI27(Fig. 1B-D, Fig. 2C), thus demonstrating immune priming. However, numerous other ISGs returned to baseline levels in the RU control but demonstrated enhanced induction after restimulation (RR) (Fig. S2C), as is indicative of immune training. Of note, one pathway that was specifically enriched for training but showed few priming effects was NF-κB (Fig. 2D, E). In contrast, priming predominantly affected antiviral ISGs (Fig. 2D,E), thus providing novel evidence for the existence of a functional state (RU) in which an acute inflammatory response is limited while direct antiviral activities are maintained. However, simultaneous immune priming and training might not be a special feature of RIG-I activation. Indeed, similar effects would be expected for specific MDA5 activation, which also signals through MAVS and TBK1(Rehwinkel & Gack, 2020). However, the lack of a selective MDA5 ligand complicates such studies. Moreover, in most published TI studies, a priming control comparable to RU is not included, and, thus, delineating transcriptional differences in priming and training is not readily possible for most stimuli at this time(Ifrim *et al*, 2014; Saeed *et al*, 2014; Quintin *et al*, 2012). Interestingly, we also observed that epithelial cells and fibroblasts showed a differential pattern of genes that remain upregulated in RU when compared to monocytes. Thus, the specific genes that are subject to priming and training may not only be stimulus-but also cell type-specific, providing a first insight into the finely-tuned complexity underlying innate immune memory responses.

Despite the established use of monocytes in the TI model(Bekkering *et al*, 2016a; Domínguez-Andrés *et al*, 2021), this cell population is short-lived, thus limiting its capability to contribute to long-term cellular memory. However, our data demonstrate that 3pRNA training is also induced in epithelial cells and fibroblasts, cells in which 3pRNA stimulation only had a limited effect on the cell cycle and cell number (Fig. S4J-M). Epithelial cells and fibroblasts, due to their anatomical location and their lifespan, may have more potential to contribute to immune training *in vivo*. Moreover, since immune training could be reproduced in rapidly dividing A549 cells (Fig. S4P-R), the preservation of TI effects across cell division can be assumed. Future research will be necessary to determine the quality and duration of such training effects in different cell types and organ systems.

Mechanistically, STAT1 signaling was found to be essential for RIG-I induced training effects (Fig. 5A-D). Moreover, in line with the requirement of STAT1, IFNβ-induced training of murine embryonic fibroblasts was previously reported (Kamada *et al*, 2018). However, in A549 only a weak training effect for IFNβ was observed, and this effect required high doses of IFNβ (100,000U/mL) (Fig. 5F-M). Thus, while type-I IFN release and subsequent STAT1 signaling was necessary for RIG-I training, it was not sufficient to recapitulate the effect of RIG-I activation in this context. It is possible that these discrepancies are due to differences in species, developmental stages or cell types, all of which merit further investigation. Moreover, while IFN signaling is clearly required for robust RIG-I expression (Schlee & Hartmann, 2016; Rehwinkel & Gack, 2020), the IFN-independent effector functions of RIG-I and other antiviral PRRs remain poorly understood(Xu *et al*, 2017; Stone & Gale, 2017), complicating the molecular elucidation of these effects. Of note, it has been shown previously that STAT1-deficient patients display defective *β*-Glucan training(Ifrim *et al*, 2015). Since *β*-Glucan/Dectin-1 signaling is molecularly quite different from RIG-I, it is striking that these two means of innate immune training utilize a common pathway. While STAT1 in RIG-I signaling acts immediately downstream of paracrine or autocrine IFNAR activation, resulting in a feedforward loop(Schlee & Hartmann, 2016; Rehwinkel & Gack, 2020), the role of STAT1 in *β*-Glucan training is less clear and has been attributed to IFNγ-signaling from T- and NK-cells(Ifrim *et al*, 2015). However, Dectin-1 signaling is also known to induce type-I IFN through IRF5(del Fresno *et al*, 2013; Mata-Martínez *et al*, 2022) which may contribute to its training effects. Furthermore, a very recent study has also shown that type I IFN-signaling is essential for LPS mediated TI in alveolar macrophages(Zahalka *et al*, 2022). Thus, type I IFN signaling may even be a universal feature of TI and should be further investigated for stimuli such as the BCG vaccine. The possibly broader role of STAT1 for diverse TI types has important clinical implications for clinical monitoring of STAT1-deficient patients(Tangye *et al*, 2022) and for patients receiving JAK-STAT inhibitors, which are commonly used in oncologic and rheumatologic treatment regimens(Tanaka *et al*, 2022; Hosseini *et al*, 2020).

RIG-I training also resulted in functional consequences for the innate immune response. Based on the results of 3’mRNAseq, we investigated the effect of 3pRNA training on P2RX7 signaling. Trained cells demonstrated P2RX7 sensitization, enabling activation and thereby the release of IL-1β at lower levels of extracellular ATP than untreated cells (Fig. 2H-J). The standard dose of extracellular ATP used to activate P2RX7 and the inflammasome is 5 mM, which is close to the total intracellular concentration of ATP in most cell types(Greiner & Glonek, 2021), and thus clearly supraphysiological in the extracellular space. RIG-I training substantially reduced the amount of ATP necessary for NLRP3 inflammasome activation and obviated the need for previous TLR2- or TLR4-mediated priming (Fig. S2S), rendering P2X7-mediated inflammasome activation more feasible in a physiological context. RIG-I agonists have been employed in numerous antiviral and antitumoral therapies(Corp., 2018; Bek *et al*, 2019; Yong & Luo, 2018; Medicine) in which P2RX7 activation may play a role. Indeed, P2RX7 activation has been previously shown to be beneficial in antiviral settings by activating the NLRP3 inflammasome and by inducing IFNβ expression(Zhang *et al*, 2017). Moreover, P2RX7 activation has been shown to contribute to antitumor immunity(Ghiringhelli *et al*, 2009), although its role in this context is controversial(Hreich *et al*, 2021), and a P2RX7-activating small molecule was recently shown to improve antitumor immune responses as well as the response to checkpoint immunotherapy(Laetitia *et al*, 2021). Notably, type-I IFN signaling, as is robustly induced by RIG-I activation, is generally considered a negative regulator of inflammasome activation(Reboldi *et al*, 2014; Guarda *et al*, 2011a). Thus, the potential contribution of this unexpected RIG-I-mediated P2RX7 induction to the antiviral and antitumoral effects of RIG-I agonists awaits further mechanistic and functional characterization.

The described effects of 3pRNA-mediated training are not only relevant for the understanding of TI mechanisms but also have clinical implications considering dosage and intervals of RIG-I agonist administration for antiviral and antitumor therapy. Particularly, prophylactic settings require the administration of RIG-I agonists for extended periods(Marx *et al*, 2022). In our study, we compared the effect of single and repetitive stimulation on antiviral protection against IAV, a negative-strand RNA virus, and against CMV, a DNA virus, *in vitro* (Fig. 1I,J). Consistent with what has been observed for other RNA and DNA viruses, including RSV and HSV-1(Liu *et al*, 2016; Schwab *et al*, 2022a), 3pRNA stimulation 24h prior to infection protected monocytes against both IAV and CMV infection. In the CMV model, single 3pRNA stimulation at day 0 significantly reduced infection rate at day 7 indicating unspecific antiviral protection upon 3pRNA training. Similar but less pronounced effects for single stimulation (UR, RU) were found after infection of A549 cells (Fig. 4L,M Fig. S4Z). Moreover, repetitive stimulation (RR) did not weaken the antiviral effect of RIG-I agonist application (Fig.1I,J). Indeed, in A549 cells repetitive stimulation induced the strongest antiviral response to CMV, indicating a potentiation of the antiviral effects of RIG-I with serial application. For the other conditions (Fig. 1I,J, Fig. 4Ls), any additive effects of serial stimulation are likely obscured by the potent antiviral response to single stimulation. Although our data clearly demonstrate that tolerance to RIG-I activation does not occur in this context, they also indicate that RIG-I priming and training effects may necessitate the adaptation of the RIG-I agonist dose in order to minimize potential side effects while optimizing therapeutic efficacy.

Altogether, our study uses a selective RIG-I agonist to demonstrate the potential of specific cytosolic nucleic-acid immune sensing to induce innate immune priming and training, adding RIG-I activation to an arsenal of other agents, notably *β*-Glucan and BCG, which can be employed for non-antigen specific immune protection. Future studies will be necessary to explore the effects of RIG-I-induced TI *in vivo*, as well as to characterize the potential effects of other cytosolic nucleic-acid receptors, such as cGAS and MDA5, on innate immune memory. These insights will not only help to elucidate the effects of single immune sensing pathways on innate immune memory but will also help to understand how more complex immune stimuli such as viruses, which activate and inhibit multiple innate immune receptors, tolerize, prime and train our innate immune response.

## MATERIALS AND METHODS

### Study Design

The primary objective of the study was to elucidate the effects of repetitive RIG-I activation in human non-malignant cells in a controlled laboratory experiment. RT-PCR, ELISA, Flow cytometry, 3’mRNA Sequencing and ATAC Sequencing were used to investigate the effects of RIG-I activation by synthetical RNA. Experiments were performed by non-blinded investigators in the laboratory of the Institute of Clinical Chemistry and Clinical Pharmacology of the university hospital Bonn. Primary endpoints were selected prospectively, performing at least 3 independent experiments for FACS, RT-PCR and ELISA.

ATAC and 3’mRNA Sequencing experiments were performed with indicated number of donors. For 3’mRNA Sequencing technical replicates and donors were included in the model. General materials are listed in table S3., Hardware used in Table S3.

If necessary, experiments were excluded if control samples showed excessive cell death or unspecific staining.

### Cell culture

Cells were cultured at 37°C and 5% CO2 and regularly tested for mycoplasma. Cell counts were determined using the TC-20 cell counter (Bio-Rad, Hercules, USA). Cells were plated using 100 μl per 96well and 1000 μl per 12 well. Cell culture reagents were obtained by Thermo Fisher Scientific (Waltham, USA) if not stated otherwise.

#### Monocytes

Buffy Coats from healthy donors were obtained from the Institute of Experimental Haematology and Transfusion Medicine, University Hospital Bonn. Peripheral blood mononuclear cells (PBMCs) were isolated by density gradient centrifugation. Isolation of CD14+ Monocytes was performed using anti-CD14 microbeads (MACS technology, Miltenyi, Radolfzell, Germany) according to the manufacturer’s instruction. Cells were plated in culture medium (RPMI 1640, 10% fetal bovine serum (FBS), Antibiotic-Antimycotic (100 IE/ml Penicillin, 100 μg/ml Streptomycin, 250 pg/ml Amphotericin B), Sodium Pyruvate, MEM non-essential amino acids,) containing 100 pg/ml M-CSF (ImmunoTools, Friesoythe, Germany) at a density of 10^6^ cells/ml. Media was exchanged at day 1 and day 4. 2 mM EDTA in PBS was used for detachment of monocytes.

#### Primary epithelial cells

Epithelial cells were isolated from human adenoids as described (Michea *et al*, 2013), and cultivated in Keratinocyte Growth Medium 2 (Heidelberg, Germany) supplemented with Antibiotic-Antimycotic and adjusted CaCl2 concentration (0.1 mM). For passaging, cells were detached using 0.05% Trypsin for 5 min at 37 °C and 5% CO2, which was subsequently neutralized with RPMI 1640 supplemented with 10% FBS and removed by centrifugation. For experiments cells were plated at 1.2*10^5^ cells/ml density the day before the experiment.

#### Fibroblasts

Fibroblasts were isolated as described above for epithelial cells using supplemented DMEM (10% FBS, 100 IE/ml Penicillin, 100 μg/ml Streptomycin) as culture medium. Medium was exchanged every 3-4 days. For passaging, cells were detached using 0.05% Trypsin for 5 min at 37°C 5% CO2, which was subsequently neutralized with DMEM. For experiments cells were plated at 2*10^5^ cells/ml density the day before the experiment.

#### A549 cells

Wildtype and *STAT1^-/-^* A549 were cultured in DMEM (10% FBS, 100 IE/ml penicillin, 100 μg/ml streptomycin) using 0.05% trypsin for passaging. For experiments cells were plated at 1.6*10^5^ cells/ml density the day before the experiment. 24 hours after the first stimulation cells were detached and cultivated in a flask until day 5 when cells were plated for stimulation on day 6.

### Stimulation

Cells were stimulated on day 0 and day 6 (d0 and d6) using synthetic RNA (25 ng/ml for monocytes, 100 ng/ml for fibroblasts, epithelial cells and A549) which was transfected according to the manufacturer’s instructions using 0.1 μl (100 ng/ml RNA) or 0.03 μl (25 ng/ml RNA) Lipofectamine 2000 per 96 well. Medium was exchanged 24h after the stimulation.

For RIG-I activation the following synthetically produced triphosphorylated double stranded RNA (3pRNA, R) was used: sense 5’pppGACGCUGACCCUGAAGUUCAUCUU3’, antisense: 5’AAGAUGAACUUCAGGGUCAGCGUC 3’. Synthetic RNA was obtained from Biomers (Ulm, Germany), 5’-triphosphate synthesis was performed in-house as previously described(Goldeck *et al*, 2014). Non-stimulatory RNA CACACACACACACACACACAC (C) or 3pRNA-antisense (A) was used as a negative transfection control. For interferon-*β* stimulation, interferon was added at the indicated concentrations (1000 U/ml-100,000 U/ml final concentration, ImmunoTools, Friesoythe, Germany). Medium was exchanged 24h after d0-stimulation. For inflammasome activation, d6-monocytes were stimulated with either 3pRNA as described above or 1 μg/ml P3CSK (InvivoGen, San Diego, USA) 5h prior to ATP challenge. pH-adjusted ATP (InvivoGen, San Diego, USA) was added using the indicated final concentrations (0.2 mM - 5 mM) and supernatants were taken 3h after stimulation. The P2RX7 receptor antagonist A7400003 (5 μM, Santa Cruz Biotechnology, Santa Cruz, USA) was added 1h before ATP challenge. Nigericin (5 μM, InvivoGen, San Diego, USA) was used as a positive control for NLRP3 inflammasome activation(Watanabe *et al*, 1998). TNF (200 ng/ml, ImmunoTools, Friesoythe, Germany) and LPS (100 ng/ml, Sigma Aldrich, St. Louis, USA) were used as control stimuli for gene induction.

### Viral infection

After RNA stimulation and medium exchange on day 0 and day 1, cells were counted and replated on d5 with a density of 5*10^4^/ml (A549) and 2*10^5^/ml (monocytes). 24h after d6-stimulation cells were infected with Influenza A Virus (IAV) H1N1 pandemic strain 2009 (A/Perth/265/2009) with a multiplicity of infection (MOI) of 10 using serum-free cell culture medium. Medium was exchanged after 1h, and virus infection was quantified by FACS (details in flow cytometry section). For Cytomegalovirus (CMV) infection, cells were infected at MOI of 1 24h after d6-stimulation with HCMV TB40/E-UL122/123-mNeonGreen(CMV) expressing mNeonGreen fluorescent protein under immediate early promoter(Kasmapour *et al*, 2017). Virus infection was quantified by FACS 24h pi.

### Cytokine measurement and normalization to cell number

The cytokines CXCL10, IL6, IL1β were measured by ELISA in cell-free supernatants of stimulated cells harvested at the time indicated. ELISA assays were performed according to the manufacturer’s instructions (BD, Franklin Lakes, USA). Cell numbers were determined by quantification of DAPI-stained nuclei (5 μg/ml in PBS for 10 minutes, Thermo Fisher Scientific, Waltham, USA) after fixation of cells with 4% Formaldehyde (10 minutes at RT, Sigma Aldrich, St. Louis, USA). Images were acquired on a Cytation3 imaging reader (BioTek, Winoski, USA) and quantified using the cell-count module of BioTek Gen5 software. Cytokine levels were normalized for the number of cells prior to stimulation on day 6 (concentration/mean of counts* mean of counts in unstimulated control). Cell normalization was not performed for A549 as cells were plated at the same density before restimulation.

### Real-time PCR (RT-PCR)

Total RNA was extracted using the RNeasy mini kit (Qiagen, Hilden, Germany) modified as described in Ostendorf et. al.(Ostendorf *et al*, 2020) and cDNA was generated using Revertaid Reverse Transcriptase (Thermo Fisher Scientific, Waltham, USA) and random hexamers (Integrated DNA Technologies, Coralville, USA). RT-PCR was performed using EvaGreen Mix II (Biobudget, Krefeld, Germany), primers are listed in Sup. table 6 (Integrated DNA Technologies, Coralville, USA). CT values were normalized to GAPDH (ΔCT).

### 3’mRNA Sequencing

3’mRNA sequencing of isolated RNA was performed at the NGS-core facility Bonn. Lexogen (Vienna, Austria) QuantSeq 3’-mRNA Library Prep was used for library preparation. Sequencing was performed with Illumina (San Diego, USA) HiSeq 2500 V4 - High Output Modus with an average of 10M raw reads (50bp). After trimming of Illumina Universal Adapter with cutadapt(Marcel) and polyA trimming reads were aligned to the human genome (hg38) with STAR(Dobin *et al*, 2012). FeatureCounts(Liao *et al*, 2013) was used to assign reads to genes based on the definitions of ENSEMBL. Inclusion criteria for counting were as follows: uniquely mapped, matching strand and overlap with only one gene, i.e., non-ambiguous assignment to a single gene. Statistical analysis was performed in R environment(Team, 2020) with the R-package limma(Ritchie *et al*, 2015) taking technical and biological replicates into consideration. The Benjamini-Hochberg method was used to calculate multiple testing adjusted p-values. P-adjusted <0.05 (P-adj) was considered differentially expressed. Data visualization, such as volcano plots and heatmaps with hierarchical clustering, were generated upon logarithmic counts per million (lcpm) using R-packages ggplot2 and Pheatmap(Kolde, 2019) respectively. Hierarchical gene clustering was performed using ward D2 method. Functional enrichment analysis using differentially expressed genes (DEG) from annotated contrasts or gene clusters as input was performed using the R-packages Pathview(Luo *et al*, 2013) and ClusterProfiler(Yu *et al*, 2012). For transcription factor (TF) enrichment analysis up- and downregulated DEG from RR_UR were used as input in CheaA3(Keenan *et al*, 2019) using the ENCODE library(Consortium, 2011).

### ATAC Sequencing

3pRNA-stimulated (day 0) and unstimulated monocytes were collected at day 1 and day 6. Cells were treated with DNase detached and cryopreserved for transport to Active Motif. ATAC Sequencing was performed by Active motif (Carlsbad, USA) using Nextera Library Prep Kit for tagmentation, MinElute PCR purification Kit for purification before amplification with Agencourt AMPure SPRI beads (Beckman Coulter, Brea, USA). After Quantification with KAPA Library Quantification Kit for Illumina platforms (KAPA Biosystems, Wilmington, USA) PE42 sequencing was performed on the NextSeq 500 sequencer (Illumina, San Diego, USA). Alignment was performed with BWA algorithm(Li & Durbin, 2010). Duplicates were removed and matching pairs of uniquely mapped reads were used for further analysis. Alignments were extended in silico at their 3’-ends to a length of 200 bp and assigned to 32-nt bins along the genome.

#### Peak calling algorithm

Peaks were identified using the MACS 2.1.0 algorithm(Zhang *et al*, 2008) at a cutoff of p-value 1e-7, without control file, and with the–nomodel option. Peaks that were on the ENCODE blacklist of known false ChIP-Seq peaks were removed. For differential analysis, reads were counted in all merged peak regions (using Subread), and the replicates for each condition were compared using DESeq2 (Love *et al*, 2014).

#### Sliding window algorithm

Csaw(Lun & Smyth, 2016) was used to detect differentially bound (DB) regions. Reads with low quality and low read depth were filtered out. A negative binomial generalized log-linear model (edgeR(Robinson *et al*, 2010)) was used to fit read counts for each window and compare untreated and treated group for each day separately. FDR <0.05 was considered differentially accessible. Transcription factor motif enrichment was determined by SEA(Bailey & Grant, 2021) using differentially bound up- and downregulated regions as input.

### Flow Cytometry Analysis

Flow cytometric analysis was performed using the Attune NxT Flow Cytometer (Thermo Fisher Scientific, Waltham, USA) and later analyzed with FlowJo (Tree Star) for all readouts. Debris, duplicates and live-dead staining positive cells were excluded before downstream analysis. If not otherwise stated, reagents were obtained by Thermo Fisher Scientific (Waltham, USA).

#### Extracellular staining

Cells were stimulated for 18h with 3pRNA before detachment with TrypLE for 5minutes. After live-dead staining (Fixable Viability Dye eFluor 780 1:1000 in PBS, 65-0865-14), cells were stained in FACS Buffer (1:200, PBS, 2% FBS, 2 mM EDTA, 0,05% NaN3) with antibodies recognizing: CD14 PECy7 (555399, BD Biosciences, Franklin Lakes, USA), CD16 BV510 (563830, BD Biosciences, Franklin Lakes, USA), CD40 APC (555591, BD Biosciences, Franklin Lakes, USA), CD69 BV650 (310934, BioLegend, San Diego, USA), CD80 PE (340294, BD Biosciences, Franklin Lakes, USA), CD83 FITC (11-0839-41, Thermo Fisher Scientific, Waltham, USA)

#### Intracellular CXCL10 staining

Cells were stimulated with 3pRNA for 8h and 18h as described above. GolgiStop (BD Biosciences, Franklin Lakes, USA) was added 5h before detachment of cells with TrypLE for 5minutes. Cells were stained with Fixable Viability Dye eFluor 506 (1:1000 in PBS, 65-0866-14) and later CD14-APC (555399, BD Biosciences, Franklin Lakes, USA) and CD16-FITC (555406, BD Biosciences, Franklin Lakes, USA) 1:200 in FACS Buffer) with FC-Block (564220, BD Biosciences, Franklin Lakes, USA).

CXCL10 (519504, BioLegend, San Diego, USA) staining was performed using the FOXP3 Kit (00-5523-00) according to the manufacturer’s instructions, followed by FC-Block and incubation of the cells for 10min at the last washing step with permeabilization buffer. PE-Isotype was used as negative control for unspecific staining.

#### DAPI uptake kinetics

Monocytes were detached using TrypLE after repetitive stimulation and resuspended in prewarmed DAPI-uptake buffer (140 mM Sucrose, 5 mM KCl, 20 mM HEPES, 5 mM D-glucose, 0.1 % bovine serum albumin, 1 μg/ml DAPI) with or without P2RX7 receptor antagonist A7400003 (5 μM, Santa Cruz Biotechnologies, Santa Cruz, USA). The synthetic P2RX7 agonist BzATP (40 μM, Sigma Aldrich, St. Louis, USA) was added directly before measurement. Uptake-kinetics were determined using the median of the 90 % percentile (DAPI VL1) after excluding duplicates, without additional live-dead staining.

#### Quantification of viral infection

Cells were detached with 0.05% trypsin (A549) or 2 mM EDTA in PBS (Monocytes) at 37°C, stained for 5 minutes on ice with eBioscience Fixable Viability Dye eFluor 780 (1:4000 in PBS) and fixed for 20 minutes with 4% formaldehyde before analysis. For quantification of influenza A virus (IAV)-infected cells, cells were permeabilized with 0.5% Triton X100 (Sigma Aldrich, St. Louis, USA) in PBS for 10min before staining with anti-IAV NP FITC for 30 minutes (1:100 in 0.25% Triton X100, 1% FCS, 1mM EDTA, PBS). For monocytes, FC-Block was added 10 min. before staining. CMV infected cells were quantified by mNeonGreen expression.

#### Statistical analysis

Graphs show the mean and standard error if not otherwise stated. Statistical analysis was performed using two-sided ratio paired t-test and one-way ANOVA with Holm-Šídák correction for normal and lognormal distribution. Wilcoxon matched-pairs signed rank test and Friedman test with Dunn’s multiple comparison were used as non-parametric tests. Prism 9 software (GraphPad, California, USA) was used for analyses and visualization, except for sequencing data (RStudio).

The Manuscript was created using Microsoft Office (Redmond, USA) and Adobe Illustrator (San Jose, USA). Biorender (Toronto, Canada) was used for graphics.

## Acknowledgments

We thank Meghan Campbell for her critical reading of this manuscript and Katrin Reiners for helpful scientific discussion.

## Author contributions

Conceptualization: MA, JvdB, GH, EB

Methodology: MA, SN, AB, PR, LC-S, MS, GH, EB

Software: MA, SN, AB, TSG

Validation: KB, SSch, SSol, EB

Formal Analysis: MA, SN, AB, EB

Investigation: MA, KB, SSch, SSol, TZ, KA-C, SM, SL, KF, EH

Resources: SH, PR, FK, LCS, MS, EH

Writing – original draft: MA, TZ, KP, GH, EB

Writing – review & editing: All authors.

Visualization: MA, SN, AB, GH, EB

Project administration: GH, EB

Supervision: MA, AB, TZ, SM, SL, LC-S, JvdB, GH, EB

Funding acquisition: GH, EB

## Data and materials availability

Additional data (e.g. Raw Sequencing data) will be made available for reviewers if requested and upon acceptance.

## Supplementary Figures

**Figure S1.** Monocytes were either unstimulated (U) or stimulated with 3pRNA (R) or control RNA (C) at day 0 (first letter) and day 6 (second letter). (A) *CXLC10* expression was determined by RT-PCR 6h after stimulation. Graph shows the mean ± SEM of -ΔCt (*GAPDH* normalized, n=6). (B) Representative image of gating strategy used for experiment shown in Fig. 1E. (C) Supernatants were collected 24h after stimulation at day 6. CXCL10 was quantified by ELISA, normalized for cell number, and plotted as the mean ± SEM of individual donors (n=10). (D) Cell numbers were determined 24h after stimulation at day 6 by quantification of DAPI-stained nuclei. Mean ± SEM (n=10) for individual donors is shown. (E) Representative images of DAPI-stained monocytes on day 6 in bright field and DAPI channel are shown for UU and RU. (F) Supernatants were collected 24h after stimulation at day 6. CXCL10 was quantified by ELISA, normalized for cell number, and plotted as the mean ± SEM of individual donors (n=5). (G) CD83 expression (mean ± SEM, n=10) was measured by FACS 18h after stimulation at day 6. Median fluorescence intensity relative to UU is shown for individual donors. (H) Representative image of gating strategy used for surface protein staining (Fig. 1F-H, Fig S1G) is shown. (J-K) Monocytes were infected with Influenza A virus (IAV, I) or Cytomegalovirus (CMV, J) 24h after d6-stimulation. Infection was quantified by flow cytometry 8h (IAV) and 24h (CMV) post infection. Uninfected control samples not shown in Fig. 1 I,J are plotted as mean ± SEM of individual donors. (K) Representative image of gating strategy for quantification of virus infection is shown. P-values were determined by one-way ANOVA using Holm-Šídák correction (A,C,F), Friedmann test with Dunn’s correction (D) and Wilcoxon matched-pairs signed rank test (G). Ns = not significant; *P ≤ 0.05, **P≤ 0.01, ***P ≤ 0.001, ****P ≤ 0.000

**Figure S2.** RNA for 3’mRNA sequencing was isolated from monocytes (n=4) harvested 24h after stimulation at day 0 and 6h after stimulation at day 6. (A) Principal component analysis for mRNA sequencing results is shown. (B) Total number of significantly up-(logarithmic fold-change, LFC >0) and downregulated (LFC <0) genes (total = differentially expressed genes, DEG) are shown for the indicated comparisons. (C) Venn Diagram shows overlap of DEG for indicated contrasts. (D) Logarithmic foldchanges (LFC) of single versus repetitive stimulation (RR_UU vs. UR_UU) are plotted. Correlation was analyzed using Spearman correlation analysis and linear regression modeling. (E) Heatmap shows the most significant DEG for RR_UR. (F) The most enriched GO terms determined using DEG for the indicated comparisons as input are shown. (G) LFC for single (UR_UU) and repetitive stimulation (RR_UU) are shown for tolerized genes (LFC>0.5, P-adj <0.05 in UR_UU and LFC_UR > LFC_RR). (H) Four main clusters were identified by expression similarity clustering using wardD2 for samples isolated at day 6. Treatment-dependent gene expression and top-enriched KEGG pathways are shown for each cluster. (I) Heatmap shows most significant genes in RR_UR out of genes listed in KEGG NOD-like receptor signaling pathway. (J-M) Gene expression for *IL1B* (J), *NLRP3* (K), *P2RX7* (L) and *IFIT1* (M) was determined using RT-PCR 8h after stimulation with the indicated stimuli. Graphs show the −ΔCt ± SEM of individual donors (n=5, *GAPDH* normalized) with P-values determined by one-way ANOVA using Holm-Šidák correction. (N-Q) Gene expression (Log2 counts per million, LCPM) for *IL1B* (N), *NLRP3* (O), *P2RX7* (P) and *IFIT1* (Q) is shown for indicates samples. (R) Representative image of gating strategy for DAPI-Uptake (Fig. 2 H,I) is shown. (S,T) Controls not shown in Fig. 2J. Pam3Cys stimulation was used as a priming control (UP). Graph is plotted as the mean ± SEM of individual donors (n=8). (U) Heatmap shows LFC of selected pattern recognition receptors. Not significantly regulated genes are shown in grey, not expressed genes are marked with *. Ns = not significant; *P ≤ 0.05, **P≤ 0.01, ***P≤ 0.01, ***P≤ 0.001.

**Figure S3.** (A) Schematic overview of the experimental setting. Day 0 3pRNA stimulated (R) and unstimulated (U) monocytes were harvested at day 1 and at day 6 for ATAC sequencing (n=2). (B-C) Principal component analysis was performed using differentially accessible regions (DAR) determined by peak-calling algorithm (B) and sliding-window approach (C). (D) The amount of DAR comparing RU vs. UU at day 6 is shown for the sliding-window and peak-calling approaches. (E) Clustered heatmaps show the most significant, gene-annotated, DAR for RU vs. UU obtained by peak-calling algorithm. (F) Reads for individual samples at d6 are shown for *IFI27*, the DAR determined by sliding-window approach is highlighted in red. (G-I) mRNA sequencing counts (monocytes, n=4) are shown for *CCL2* (G), *IFI27* (H), and *MX1* (I) as logarithmic counts per million (LCPM). (J-K) Venn diagram shows the overlap of genes from DAR for sliding-window and peak-calling approach and differentially expressed genes (DEG) for mRNA sequencing by comparing RR to UR (J) and RU to UU (K). (L) Heatmap shows transcriptional expression of the most overrepresented transcription factor binding motifs obtained by SEA (DAR RU vs. UU). Not expressed genes are shown in grey (NA).

**Figure S4.** RNA for 3’mRNA sequencing was isolated 6h after stimulation of epithelial cells (n=3) and fibroblasts (n=4). (A-D) Logarithmic fold-changes (LFC) of single versus repetitive stimulation (RR_UU vs. UR_UU) are plotted for epithelial cells (A) and fibroblasts (B). Correlation was analyzed using Spearman correlation analysis and linear regression modeling. Significantly up- and downregulated genes (LFC >0/<0) in epithelial cells (C) and fibroblasts (D) are shown for each comparison. (E,F) The Overlap of differentially expressed genes (DEG) for the indicated contrasts is shown in epithelial cells (E) and fibroblasts (F). (G,H) KEGG Pathway enrichment analysis using DEG for the annotated contrasts in epithelial cells (G) and fibroblasts (H) is plotted. (I) Expression overlaps of all genes included in the mRNA sequencing for monocytes, epithelial cells, and fibroblasts are shown. (J,K) The most variant genes included in the KEGG-pathway cell cycle (hsa04110) are shown in the heatmap for epithelial cells (J) and fibroblasts (K).(L,M) Cell counts were determined 10h and 24h after the second stimulation in epithelial cells (L) and fibroblasts (M) by quantification of DAPI-stained nuclei and plotted as the mean ± SEM for individual donors (L n=5, M n=10). P-values were determined by Friedman test using Dunn’s correction comparing each treatment to unstimulated control (UU). (N,O) Heatmaps show the most variant genes included in the KEGG-pathway NFkappaB signaling for epithelial cells (N) and fibroblasts (O). (P-U) Interferon-stimulated gene expression (*IFIT1, RSAD2, IFI27*) in A549 cells was determined 6h after stimulation at day 6 (P-R) and day 28 (S-U) by RT-PCR. Graph shows mean -ΔCt ± SEM (n=6 P-R, n=4 S-U, *GAPDH* normalized). P-values were determined by one-way ANOVA using Holm-Šídák correction (P-R) and Friedman test using Dunn’s correction (S-U). (V) Controls not shown for IRF3 phosphorylation western blot in Fig. 4H. (W) CXCL10 production 10h after d6-stimulation was determined in monocytes and normalized for cell number. Graph shows the mean ± SEM (n=6) with P-values determined by one way ANOVA using Holm-Šídák correction. (X,Y) Heatmap show LFC of selected pattern recognition receptors for epithelial cells (X) and fibroblasts (Y). Not significantly regulated genes are shown in grey, not expressed genes are marked with *. (Z) Representative image of gating strategy used for quantification of virus infection in Figure 4 L,M is shown. Ns = not significant, *P≤0.05, **P≤0.01, ***P≤0.001, ****P≤0.0001

**Figure S5.** (A) β-Actin and STAT1 determined by western blot are shown for A549 wildtype (WT) and STAT1^-/-^ (KO) cells. (B) Type-I interferon (IFN) release 24h after primary 3pRNA stimulation was quantified by using HEK-Blue type-I IFN reporter cells. Boxplot shows minimum and maximum as whiskers (n=11). (C-F) RIG-I signaling was determined by RT-PCR 6h after A549 WT cells were challenged with 100,000 U/ml IFNβ (I) or 3pRNA (R) at day 0 and/or day 6. Graphs show the mean −ΔCt ± SEM (n=6, *GAPDH* normalized) for indicated genes with P-values determined by one-way ANOVA using Holm-Šídák correction. (G-J) RIG-I signaling was determined by RT-PCR 6h after A549 WT cells were challenged with 2000 U/ml IFNβ (I) or 3pRNA (R) at day 0 and/or day 6. Graphs show the mean −ΔCt ± SEM (n=3, *GAPDH* normalized) for indicated genes. ns = not significant, *p ≤ 0.05, **p ≤ 0.01, ***p ≤ 0.01

## Supplementary Materials

Table S1: Results obtained by SEA transcription factor motif enrichment analysis using differentially accessible regions (DAR) obtained by sliding window approach comparing RU vs. UU as input (upregulated DAR LFC >0, downregulated DAR LFC<0).

Table S2: Results obtained by ChEA3 transcription factor overrepresentation analysis (ENCODE dataset) using differentially expressed genes in RR vs. UR (mRNA Sequencing, monocytes) as input (upregulated DEG LFC >0, downregulated DEG LFC<0).

Table S3: List of Reagents used

Table S4: List of Hardware used

Table S5: List of RT PCR Primers used

## References and Notes

Ablasser A, Poeck H, Anz D, Berger M, Schlee M, Kim S, Bourquin C, Goutagny N, Jiang Z, Fitzgerald KA, et al (2009) Selection of molecular structure and delivery of RNA oligonucleotides to activate TLR7 versus TLR8 and to induce high amounts of IL-12p70 in primary human monocytes. Journal of immunology 182: 6824–33

Arts RJW, Joosten LAB & Netea MG (2018a) The Potential Role of Trained Immunity in Autoimmune and Autoinflammatory Disorders. Frontiers in Immunology 9

Arts RJW, Moorlag SJCFM, Novakovic B, Li Y, Wang S-Y, Oosting M, Kumar V, Xavier RJ, Wijmenga C, Joosten LAB, et al (2018b) BCG Vaccination Protects against Experimental Viral Infection in Humans through the Induction of Cytokines Associated with Trained Immunity. Cell Host Microbe 23: 89–100.e5

Bailey TL & Grant CE (2021) SEA: Simple Enrichment Analysis of motifs. Biorxiv: 2021.08.23.457422

Bartok E & Hartmann G (2020a) Immune Sensing Mechanisms that Discriminate Self from Altered Self and Foreign Nucleic Acids. Immunity 53: 54–77

Bauernfeind FG, Horvath G, Stutz A, Alnemri ES, MacDonald K, Speert D, Fernandes-Alnemri T, Wu J, Monks BG, Fitzgerald KA, et al (2009) Cutting edge: NF-kappaB activating pattern recognition and cytokine receptors license NLRP3 inflammasome activation by regulating NLRP3 expression. Journal of immunology (Baltimore, Md: 1950) 183: 787–791

Bek S, Stritzke F, Wintges A, Nedelko T, Böhmer DFR, Fischer JC, Haas T, Poeck H & Heidegger S (2019) Targeting intrinsic RIG-I signaling turns melanoma cells into type I interferon-releasing cellular antitumor vaccines. Oncoimmunology 8: e1570779

Bekkering S, Blok BA, Joosten LAB, Riksen NP, Crevel R van & Netea MG (2016a) In Vitro Experimental Model of Trained Innate Immunity in Human Primary Monocytes. Clin Vaccine Immunol 23: 926–933

Bekkering S, Blook BA, Joosten LAB, Riksen NP, Crevel R van, Netea MG & Rosenberg HF (2016b) In Vitro Experimental Model of Trained Innate Immunity in Human Primary Monocytes. Clin Vaccine Immunol 23: 926–933

Bergstrøm B, Aune MH, Awuh JA, Kojen JF, Blix KJ, Ryan L, Flo TH, Mollnes TE, Espevik T & Stenvik J (2015) TLR8 Senses Staphylococcus aureus RNA in Human Primary Monocytes and Macrophages and Induces IFN-β Production via a TAK1-IKKβ-IRF5 Signaling Pathway. J Immunol 195: 1100–1111

Berthelot J-M & Sibilia J (2019) Trained Immunity and Autoimmune Disease: Did Eve Sin before Adam? Joint Bone Spine 86: 293–295

Butcher SK, O’Carroll CE, Wells CA & Carmody RJ (2018) Toll-Like Receptors Drive Specific Patterns of Tolerance and Training on Restimulation of Macrophages. Front Immunol 9: 933

Cheemarla NR, Watkins TA, Mihaylova VT, Wang B, Zhao D, Wang G, Landry ML & Foxman EF (2021) Dynamic innate immune response determines susceptibility to SARS-CoV-2 infection and early replication kinetics. Journal of Experimental Medicine 218

Cirovic B, Bree LCJ de, Groh L, Blok BA, Chan J, Velden WJFM van der, Bremmers MEJ, Crevel R van, Händler K, Picelli S, et al (2020) BCG Vaccination in Humans Elicits Trained Immunity via the Hematopoietic Progenitor Compartment. Cell Host Microbe 28: 322–334.e5

Clark IA, Allison AC & Cox FE (1976) Protection of mice against Babesia, and Plasmodium with BCG. Nature 259: 309–311

Coch C, Stümpel JP, Lilien-Waldau V, Wohlleber D, Kümmerer BM, Bekeredjian-Ding I, Kochs G, Garbi N, Herberhold S, Schuberth-Wagner C, et al (2017a) RIG-I Activation Protects and Rescues from Lethal Influenza Virus Infection and Bacterial Superinfection. Mol Ther 25: 2093–2103

Coch C, Stumpel JP, Lilien-Waldau V, Wohlleber D, Kummerer BM, Bekeredjian-Ding I, Kochs G, Garbi N, Herberhold S, Schuberth-Wagner C, et al (2017b) RIG-I Activation Protects and Rescues from Lethal Influenza Virus Infection and Bacterial Superinfection. Molecular therapy: the journal of the American Society of Gene Therapy 25: 2093–2103

Consortium TEP (2011) A User’s Guide to the Encyclopedia of DNA Elements (ENCODE). Plos Biol 9: 1–21

Corp. MS&D (2018) intratumoral/intralesional administration of mk-4621/jetpei^TM^ with or without pembrolizumab in participants with advanced/metastatic or recurrent solid tumors (mk-4621-002).

Dang EV, McDonald JG, Russell DW & Cyster JG (2017) Oxysterol Restraint of Cholesterol Synthesis Prevents AIM2 Inflammasome Activation. Cell 171: 1057–1071.e11

Decker T (2021) The early interferon catches the SARS-CoV-2. J Exp Medicine 218: e20211667

del Fresno C, Soulat D, Roth S, Blazek K, Udalova I, Sancho D, Ruland J & Ardavín C (2013) Interferon-beta; Production via Dectin-1-Syk-IRF5 Signaling in Dendritic Cells Is Crucial for Immunity to C.albicans. Immunity 38: 1176–1186

Divangahi M, Aaby P, Khader SA, Barreiro LB, Bekkering S, Chavakis T, Crevel R van, Curtis N, DiNardo AR, Dominguez-Andres J, et al (2021a) Trained immunity, tolerance, priming and differentiation: distinct immunological processes. Nat Immunol 22: 2–6

Divangahi M, Aaby P, Khader SA, Barreiro LB, Bekkering S, Chavakis T, Crevel R van, Curtis N, DiNardo AR, Dominguez-Andres J, et al (2021b) Trained immunity, tolerance, priming and differentiation: distinct immunological processes. Nat Immunol 22: 2–6

Dobin A, Davis CA, Schlesinger F, Drenkow J, Zaleski C, Jha S, Batut P, Chaisson M & Gingeras TR (2012) STAR: ultrafast universal RNA-seq aligner. Bioinformatics 29: 15–21

Domínguez-Andrés J, Arts RJW, Bekkering S, Bahrar H, Blok BA, Bree LCJ de, Bruno M, Bulut Ö, Debisarun PA, Dijkstra H, et al (2021) In vitro induction of trained immunity in adherent human monocytes. Star Protoc 2: 100365

Fanucchi S, Domínguez-Andrés J, Joosten LAB, Netea MG & Mhlanga MM (2020) The Intersection of Epigenetics and Metabolism in Trained Immunity. Immunity 54: 32–43

Faustman DL & Davis M (2015) Historic and Emerging Applications of BCG in Prevention and Treatment of Disease. Medical Res Archives 2

Feld M & Schuberth-Wagner C (2022) Method for designing RIG-I ligands.

Feng Y, R. P David, J. C David & C. W Nicholas (2020) From reads to insight: a hitchhiker’s guide to ATAC-seq data analysis. Genome Biol 21: 22

Freudenberg MA & Galanos C (1988) Induction of tolerance to lipopolysaccharide (LPS)-D-galactosamine lethality by pretreatment with LPS is mediated by macrophages. Infect Immun 56: 1352–1357

Fung S-Y, Siu K-L, Lin H, Chan C-P, Yeung ML & Jin D-Y (2022) SARS-CoV-2 NSP13 helicase suppresses interferon signaling by perturbing JAK1 phosphorylation of STAT1. 12: 36

Garcia-Valtanen P, Guzman-Genuino RM, Williams DL, Hayball JD & Diener KR (2017) Evaluation of trained immunity by beta-1,3(d)-glucan on murine monocytes in vitro and duration of response in vivo. Nature Publishing Group 95: 601–610

Garly M-L, Martins CL, Balé C, Baldé MA, Hedegaard KL, Gustafson P, Lisse IM, Whittle HC & Aaby P (2003) BCG scar and positive tuberculin reaction associated with reduced child mortality in West Africa A non-specific beneficial effect of BCG? Vaccine 21: 2782–2790

Ghiringhelli F, Apetoh L, Tesniere A, Aymeric L, Ma Y, Ortiz C, Vermaelen K, Panaretakis T, Mignot G, Ullrich E, et al (2009) Activation of the NLRP3 inflammasome in dendritic cells induces IL-1β-dependent adaptive immunity against tumors. Nat Med 15: 1170–1178

Giard DJ, Aaronson SA, Todaro GJ, Arnstein P, Kersey JH, Dosik H & Parks WP (1973) In vitro cultivation of human tumors: establishment of cell lines derived from a series of solid tumors. J Natl Cancer I 51: 1417–23

Goldeck M, Tuschl T, Hartmann G & Ludwig J (2014) Efficient Solid-Phase Synthesis of pppRNA by Using Product-Specific Labeling - Goldeck - 2014 - Angewandte Chemie - Wiley Online Library.

Greiner JV & Glonek T (2021) Intracellular ATP Concentration and Implication for Cellular Evolution. Biology 10: 1166

Guarda G, Braun M, Staehli F, Tardivel A, Mattmann C, Förster I, Farlik M, Decker T, Du Pasquier RA, Romero P, et al (2011a) Type I Interferon Inhibits Interleukin-1 Production and Inflammasome Activation. Immunity 34: 213–223

Guarda G, Braun M, Staehli F, Tardivel A, Mattmann C, Förster I, Farlik M, Decker T, Pasquier RAD, Romero P, et al (2011b) Type I interferon inhibits interleukin-1 production and inflammasome activation. Immunity 34: 213–223

Guillon A, Arafa EI, Barker KA, Belkina AC, Martin IMC, Shenoy AT, Wooten AK, Ana CLD, Dai A, Labadorf A, et al (2020) Pneumonia recovery reprograms the alveolar macrophage pool. Jci Insight 5

Gyssens IC & Netea MG (2019) Heterologous effects of vaccination and trained immunity. Clin Microbiol Infec 25: 1457–1458

Hartmann G (2017) Nucleic Acid Immunity. Adv Immunol 133: 121–169

Hoesel B & Schmid JA (2013) The complexity of NF-κB signaling in inflammation and cancer. Mol Cancer 12: 86

Hosseini A, Gharibi T, Marofi F, Javadian M, Babaloo Z & Baradaran B (2020) Janus kinase inhibitors: A therapeutic strategy for cancer and autoimmune diseases. Journal of Cellular Physiology 235: 5903–5924

Hreich SJ dit, Benzaquen J, Hofman P & Vouret-Craviari V (2021) To inhibit or to boost the ATP/P2RX7 pathway to fight cancer—that is the question. Purinerg Signal 17: 619–631

Ifrim DC, Quintin J, Joosten LAB, Jacobs C, Jansen T, Jacobs L, Gow NAR, Williams DL, Meer JWM van der, Netea MG, et al (2014a) Trained Immunity or Tolerance: Opposing Functional Programs Induced in Human Monocytes after Engagement of Various Pattern Recognition Receptors. Clin Vaccine Immunol Cvi 21: 534–545

Ifrim DC, Quintin J, Meerstein-Kessel L, Plantinga TS, Joosten LAB, Meer JWM van der, Veerdonk FL van de & Netea MG (2015a) Defective trained immunity in patients with STAT-1-dependent chronic mucocutaneaous candidiasis. Clin Exp Immunol 181: 434–440

Interferome: www.interferome.org

Intratumoral/Intralesional Administration of MK-4621/JetPEI^-^ With or Without Pembrolizumab in Participants With Advanced/Metastatic or Recurrent Solid Tumors (MK-4621-002) (2022) ClinicalTrials.gov

Ito S, Ishii KJ, Gursel M, Shirotra H, Ihata A & Klinman DM (2005) CpG Oligodeoxynucleotides Enhance Neonatal Resistance to Listeria Infection. J Immunol 174: 777–782

Junt T & Barchet W (2015) Translating nucleic acid-sensing pathways into therapies. Nat Rev Immunol 15: 529–44

Kamada R, Yang W, Zhang Y, Patel MC, Yang Y, Ouda R, Dey A, Wakabayashi Y, Sakaguchi K, Fujita T, et al (2018) Interferon stimulation creates chromatin marks and establishes transcriptional memory. Proc National Acad Sci 115: E9162–E9171

Kasmapour B, Kubsch T, Rand U, Eiz-Vesper B, Messerle M, Vondran FWR, Wiegmann B, Haverich A, Cicin-Sain L & Jung JU (2017) Myeloid Dendritic Cells Repress Human Cytomegalovirus Gene Expression and Spread by Releasing Interferon-Unrelated Soluble Antiviral Factors. Journal of Virology 92: e01138–17

Katona B & Lindskog C (2022) The Human Protein Atlas and Antibody-Based Tissue Profiling in Clinical Proteomics. Methods Mol Biology Clifton N J 2420: 191–206

Keenan AB, Torre D, Lachmann A, Leong AK, Wojciechowicz ML, Utti V, M J Kathleen, Kropiwnicki E, Wang Z & Ma’ayan A (2019) ChEA3: transcription factor enrichment analysis by orthogonal omics integration. Nucleic Acids Res 47: W212–W224

Kleinnijenhuis J, Quintin J, Preijers F, Benn CS, Joosten LAB, Jacobs C, Loenhout J van, Xavier RJ, Aaby P, Meer JWM van der, et al (2014) Long-Lasting Effects of BCG Vaccination on Both Heterologous Th1/Th17 Responses and Innate Trained Immunity. J Innate Immun 6: 152–158

Kleinnijenhuis J, Quintin J, Preijers F, Joosten LAB, Ifrim DC, Saeed S, Jacobs C, Loenhout J van, Jong D de, Stunnenberg HG, et al (2012) Bacille Calmette-Guerin induces NOD2-dependent nonspecific protection from reinfection via epigenetic reprogramming of monocytes. P Natl Acad Sci Usa 109: 17537–42

Klemm SL, Shipony Z & Greenleaf WJ (2019) Chromatin accessibility and the regulatory epigenome. Nature Reviews Genetics: 1–14

Kolde R (2019) pheatmap: Pretty Heatmaps. CRAN R Project

Laetitia D, Serena J dit H, Jonathan B, Laetitia S, Thierry J, Xavier D, Christophe D, Bernhard R, Jean K, Cecile D, et al (2021) A small-molecule P2RX7 activator promotes anti-tumor immune responses and sensitizes lung tumor to immunotherapy. 12: 653

Larsen SB (2020) Epithelial cells: liaisons of immunity. 1–16

Li H & Durbin R (2010) Fast and accurate long-read alignment with Burrows-Wheeler transform. Bioinformatics 26: 589–595

Li MMH, MacDonald MR & Rice CM (2015) To translate, or not to translate: viral and host mRNA regulation by interferon-stimulated genes. Trends Cell Biol 25: 320–329

Liao Y, Smyth GK & Shi W (2013) featureCounts: an efficient general purpose program for assigning sequence reads to genomic features. Bioinformatics 30: 923–930

Liu Y, Goulet M-L, Sze A, Hadj SB, Belgnaoui SM, Lababidi RR, Zheng C, Fritz JH, Olagnier D & Lin R (2016) RIG-I-Mediated STING Upregulation Restricts Herpes Simplex Virus 1 Infection. J Virol 90: 9406–9419

Lopez J, Mommert M, Mouton W, Pizzorno A, Brengel-Pesce K, Mezidi M, Villard M, Lina B, Richard J-C, Fassier J-B, et al (2021) Early nasal type I IFN immunity against SARS-CoV-2 is compromised in patients with autoantibodies against type I IFNs. J Exp Medicine 218: e20211211

Love MI, Huber W & Anders S (2014) Moderated estimation of fold change and dispersion for RNA-seq data with DESeq2. Genome Biol 15: 550

Lun ATL & Smyth GK (2016) csaw: a Bioconductor package for differential binding analysis of ChIP-seq data using sliding windows. Nucleic Acids Res 44: e45

Luo, Weijun, Brouwer & Cory (2013) Pathview: an R/Bioconductor package for pathway-based data integration and visualization. Bioinformatics 29: 1830–1831

Mantovani A & Netea MG (2020) Trained Innate Immunity, Epigenetics, and Covid-19. New Engl J Med 383: 1078–1080

Mao T, Israelow B, Lucas C, Vogels CBF, Gomez-Calvo ML, Fedorova O, Breban MI, Menasche BL, Dong H, Linehan M, et al (2021) A stem-loop RNA RIG-I agonist protects against acute and chronic SARS-CoV-2 infection in mice. J Exp Medicine 219: e20211818

Marcel M Cutadapt removes adapter sequences from high-throughput sequencing reads. EMBnet.journal; Vol 17, No 1: Next Generation Sequencing Data Analysis

Marques M, Ferreira AR & Ribeiro D (2018) The Interplay between Human Cytomegalovirus and Pathogen Recognition Receptor Signaling. Viruses 10

Marx S, Kümmerer BM, Grützner C, Kato H, Schlee M, Renn M, Bartok E & Hartmann G (2022) RIG-I-induced innate antiviral immunity protects mice from lethal SARS-CoV-2 infection. Molecular therapy Nucleic acids 27: 1225–1234

Mata-Martínez P, Bergón-Gutiérrez M & Fresno C del (2022) Dectin-1 Signaling Update: New Perspectives for Trained Immunity. Frontiers in Immunology 13

Medicine USNL of a study evaluating the safety, pharmacokinetics, and antiviral efficacy of SB 9200 in subjects infected with chronic HBV. ClinicalTrialsgov

Michea P, Vargas P, Donnadieu M-H, Rosemblatt M, Bono MR, Duménil G & Soumelis V (2013) Epithelial control of the human pDC response to extracellular bacteria. European Journal of Immunology 43: 1264–1273

Mogensen TH (2019) IRF and STAT Transcription Factors - From Basic Biology to Roles in Infection, Protective Immunity, and Primary Immunodeficiencies. Front Immunol 9: 3047

Morales A, Eidinger D & Bruce AW (1976) Intracavitary Bacillus Calmette-guerin in the Treatment of Superficial Bladder Tumors. J Urology 116: 180–182

Muzio M, Bosisio D, Polentarutti N, D’amico G, Stoppacciaro A, Mancinelli R, Veer C van’t, Penton-Rol G, Ruco LP, Allavena P, et al (2000) Differential Expression and Regulation of Toll-Like Receptors (TLR) in Human Leukocytes: Selective Expression of TLR3 in Dendritic Cells. J Immunol 164: 5998–6004

Näslund C Résultats des experiences de vaccination par le BCG poursuivies dans le Norrbotten (Suède) (Septembre 1927-Décembre 1931). Vaccin Prev Tuberc Rapp Doc Paris Inst Pasteur

Netea MG, Domínguez-Andrés J, Barreiro LB, Chavakis T, Divangahi M, Fuchs E, Joosten LAB, Meer JWM van der, Mhlanga MM, Mulder WJM, et al (2020) Defining trained immunity and its role in health and disease. Nat Rev Immunol 20: 375–388

Netea MG, Joosten LAB, Latz E, Mills KHG, Natoli G, Stunnenberg HG, O’Neill LAJ & Xavier RJ (2016) Trained immunity: A program of innate immune memory in health and disease. Science 352: aaf1098

Novakovic B, Habibi E, Wang S-Y, Arts RJW, Davar R, Megchelenbrink W, Kim B, Kuznetsova T, Kox M, Zwaag J, et al (2016) β-Glucan Reverses the Epigenetic State of LPS-Induced Immunological Tolerance. Cell 167: 1354–1368.e14

Opitz B, Rejaibi A, Dauber B, Eckhard J, Vinzing M, Schmeck B, Hippenstiel S, Suttorp N & Wolff T (2007) IFNβ induction by influenza A virus is mediated by RIG-I which is regulated by the viral NS1 protein. Cellular Microbiology 9: 930–938

Ostendorf T, Zillinger T, Andryka K, Schlee-Guimaraes TM, Schmitz S, Marx S, Bayrak K, Linke R, Salgert S, Wegner J, et al (2020) Immune Sensing of Synthetic, Bacterial, and Protozoan RNA by Toll-like Receptor 8 Requires Coordinated Processing by RNase T2 and RNase 2. Immunity 52: 591–605.e6

Pedras-Vasconcelos JA, Goucher D, Puig M, Tonelli LH, Wang V, Ito S & Verthelyi D (2006) CpG Oligodeoxynucleotides Protect Newborn Mice from a Lethal Challenge with the Neurotropic Tacaribe Arenavirus. J Immunol 176: 4940–4949

Pelegrin P (2021) P2X7 receptor and the NLRP3 inflammasome: Partners in crime. Biochem Pharmacol 187: 114385

Pelegrin P & Surprenant A (2006) Pannexin-1 mediates large pore formation and interleukin-1beta release by the ATP-gated P2X7 receptor. Embo J 25: 5071–82

Pétrilli V, Papin S, Dostert C, Mayor A, Martinon F & Tschopp J (2007) Activation of the NALP3 inflammasome is triggered by low intracellular potassium concentration. Cell Death Differ 14: 1583–1589

Proietti M, Cornacchione V, Jost TR, Romagnani A, Faliti CE, Perruzza L, Rigoni R, Radaelli E, Caprioli F, Preziuso S, et al (2014) ATP-Gated Ionotropic P2X7 Receptor Controls Follicular T Helper Cell Numbers in Peyer’s Patches to Promote Host-Microbiota Mutualism. Immunity 41: 789–801

Quintin J, Saeed S, Martens JHA, Giamarellos-Bourboulis EJ, Ifrim DC, Logie C, Jacobs L, Jansen T, Kullberg B-J, Wijmenga C, et al (2012) Candida albicans Infection Affords Protection against Reinfection via Functional Reprogramming of Monocytes. Cell Host Microbe 12: 223–232

Reboldi A, Dang EV, McDonald JG, Liang G, Russell DW & Cyster JG (2014) 25-Hydroxycholesterol suppresses interleukin-1-driven inflammation downstream of type I interferon. Science 345: 679–684

Rehwinkel J & Gack MU (2020) RIG-I-like receptors: their regulation and roles in RNA sensing. Nat Rev Immunol 20: 537–551

Rehwinkel J, Tan CP, Goubau D, Schulz O, Pichlmair A, Bier K, Robb N, Vreede F, Barclay W, Fodor E, et al (2010) RIG-I Detects Viral Genomic RNA during Negative-Strand RNA Virus Infection. Cell 140: 397–408

Ribes S, Meister T, Ott M, Redlich S, Janova H, Hanisch U-K, Nessler S & Nau R (2014) Intraperitoneal prophylaxis with CpG oligodeoxynucleotides protects neutropenic mice against intracerebral Escherichia coli K1 infection. J Neuroinflamm 11: 14

Richmond JY & Hamilton LD (1969) FOOT-AND-MOUTH DISEASE VIRUS INHIBITION INDUCED IN MICE BY SYNTHETIC DOUBLE-STRANDED RNA (POLYRIBOINOSINIC AND POLYRIBOCYTIDYLIC ACIDS)*. Proc National Acad Sci 64: 81–86

Ritchie ME, Phipson B, Wu D, Hu Y, Law CW, Shi W & Smyth GK (2015) limma powers differential expression analyses for RNA-sequencing and microarray studies. Nucleic Acids Res 43: e47

Robinson MD, McCarthy DJ & Smyth GK (2010) edgeR: a Bioconductor package for differential expression analysis of digital gene expression data. Bioinformatics 26: 139–140

Rusinova I, Forster S, Yu S, Kannan A, Masse M, Cumming H, Chapman R & Hertzog PJ (2013) INTERFEROME v2.0: an updated database of annotated interferon-regulated genes. Nucleic Acids Res 41: D1040–D1046

Saeed S, Quintin J, Kerstens HHD, Rao NA, Aghajanirefah A, Matarese F, Cheng S-C, Ratter J, Berentsen K, Ent MA van der, et al (2014) Epigenetic programming of monocyte-to-macrophage differentiation and trained innate immunity. Science 345: 1251086–1251086

Sangfelt SEO (2000) Mechanisms of interferon-induced cell cycle arrest. Front Biosci 5: 479–487

Schlee M (2013) Master sensors of pathogenic RNA – RIG-I like receptors. Immunobiology 218: 1322–1335

Schlee M & Hartmann G (2016) Discriminating self from non-self in nucleic acid sensing. Nat Rev Immunol 16: 566–80

Schlee M, Roth A, Hornung V, Hagmann CA, Wimmenauer V, Barchet W, Coch C, Janke M, Mihailovic A, Wardle G, et al (2009) Recognition of 5’ triphosphate by RIG-I helicase requires short blunt double-stranded RNA as contained in panhandle of negative-strand virus. Immunity 31: 25–34

Schoggins JW & Rice CM (2011) Interferon-stimulated genes and their antiviral effector functions. Curr Opin Virol 1: 519–525

Schrum JE, Crabtree JN, Dobbs KR, Kiritsy MC, Reed GW, Gazzinelli RT, Netea MG, Kazura JW, Dent AE, Fitzgerald KA, et al (2018) Cutting Edge: Plasmodium falciparum Induces Trained Innate Immunity. J Immunol 200: 1243–1248

Schwab LSU, Farrukee R, Eléouët JF, Rameix-Welti MA, Londrigan SL, Brooks AG, Hurt AC, Coch C, Zillinger T, Hartmann G, et al (2022a) Retinoic Acid–Inducible Gene I Activation Inhibits Human Respiratory Syncytial Virus Replication in Mammalian Cells and in Mouse and Ferret Models of Infection. J Infect Dis

Schwab LSU, Londrigan SL, Brooks AG, Hurt AC, Sahu A, Deng Y-M, Moselen J, Coch C, Zillinger T, Hartmann G, et al (2022b) Induction of Interferon-Stimulated Genes Correlates with Reduced Growth of Influenza A Virus in Lungs after RIG-I Agonist Treatment of Ferrets. J Virol: e0055922

Schwab LSU, Villalón-Letelier F, Tessema MB, Londrigan SL, Brooks AG, Hurt A, Coch C, Zillinger T, Hartmann G & Reading PC (2022c) Expression of a Functional Mx1 Protein Is Essential for the Ability of RIG-I Agonist Prophylaxis to Provide Potent and Long-Lasting Protection in a Mouse Model of Influenza A Virus Infection. Viruses 14: 1547

Sender R & Milo R (2021) The distribution of cellular turnover in the human body. Nat Med 27: 45–48

Shemesh MAA (2021) SARS-CoV-2 suppresses IFNβ production mediated by NSP1, 5, 6, 15, ORF6 and ORF7b but does not suppress the effects of added interferon. PLOS Pathogens 17: 1–31

Solle M, Labasi J, Perregaux DG, Stam E, Petrushova N, Koller BH, Griffiths RJ & Gabel CA (2000) Altered cytokine production in mice lacking P2X(7) receptors. J Biological Chem 276: 125–32

Spencer JC, Ganguly R & Waldman RH (1977) Nonspecific protection of mice against influenza virus infection by local or systemic immunization with Bacille Calmette-Guérin. J Infect Dis 136: 171–5

Starr SE, Visintine AM, Tomeh MO & Nahmias AJ (1976) Effects of Immunostimulants on Resistance of Newborn Mice to Herpes Simplex Type 2 Infection. P Soc Exp Biol Med 152: 57–60

Stephen EL, Sammons ML, Pannier WL, Baron S, Spertzel RO & Levy HB (1977) Effect of a Nuclease-Resistant Derivative of PolyriboinosinicPolyribocytidylic Acid Complex on Yellow Fever in Rhesus Monkeys (Macaca mulatta). J Infect Dis 136: 122–126

Stone AEL & Gale MJ (2017) Beyond sensing: Retinoic acid inducible gene-I (RIG-I) continues to expand its antiviral effector functions. Hepatology 65: 1792–1795

Surprenant A, Rassendren F, Kawashima E, North RA & Buell G (1996) The Cytolytic P2Z Receptor for Extracellular ATP Identified as a P2X Receptor (P2X7). Science 272: 735–738

Tanaka Y, Luo Y, O’Shea JJ & Nakayamada S (2022) Janus kinase-targeting therapies in rheumatology: a mechanisms-based approach. 18: 133–145

Tangye SG, Al-Herz W, Bousfiha A, Cunningham-Rundles C, Franco JL, Holland SM, Klein C, Morio T, Oksenhendler E, Picard C, et al (2022) Human Inborn Errors of Immunity: 2022 Update on the Classification from the International Union of Immunological Societies Expert Committee. J Clin Immunol: 1–35

Team RC (2020) R: A Language and Environment for Statistical Computing.

U.S.N.L. of a study evaluating the safety, pharmacokinetics, and antiviral efficacy of SB 9200 in subjects infected with chronic HBV (2020) ClinicalTrials.gov

van den Boorn JG & Hartmann G (2013) Turning Tumors into Vaccines: Co-opting the Innate Immune System. Immunity 39: 27–37

Verthelyi D, Gursel M, Kenney RT, Lifson JD, Liu S, Mican J & Klinman DM (2003) CpG Oligodeoxynucleotides Protect Normal and SIV-Infected Macaques from Leishmania Infection. J Immunol 170: 4717–4723

Walk J, Keramati F, Bree LCJ de, Arts RJW, Blok B, Netea MG, Stunnenberg HG & Sauerwein RW (2020) Controlled Human Malaria Infection Induces Long-Term Functional Changes in Monocytes. Frontiers Mol Biosci 7: 604553

Watanabe N, Kawaguchi M & Kobayashi Y (1998) Activation of interleukin-1beta-converting enzyme by nigericin is independent of apoptosis. Cytokine 10: 645–653

Xu L, Wang W, Li Y, Zhou X, Yin Y, Wang Y, Man RA de, Laan LJW van der, Huang F, Kamar N, et al (2017) RIG-I is a key antiviral interferon-stimulated gene against hepatitis E virus regardless of interferon production. Hepatology 65: 1823–1839

Yong HY & Luo D (2018) RIG-I-Like Receptors as Novel Targets for Pan-Antivirals and Vaccine Adjuvants Against Emerging and Re-Emerging Viral Infections. Front Immunol 9: 1379

Yu G, Wang L-G, Han Y & He Q-Y (2012) clusterProfiler: an R package for comparing biological themes among gene clusters. Omics J Integr Biology 16: 284–287

Zahalka S, Starkl P, Watzenboeck ML, Farhat A, Radhouani M, Deckert F, Hladik A, Lakovits K, Oberndorfer F, Lassnig C, et al (2022) Trained immunity of alveolar macrophages requires metabolic rewiring and type 1 interferon signaling. 15: 896–907

Zhang C, He H, Wang L, Zhang N, Huang H, Xiong Q, Yan Y, Wu N, Ren H, Han H, et al (2017) Virus-Triggered ATP Release Limits Viral Replication through Facilitating IFN-β Production in a P2X7-Dependent Manner. The Journal of Immunology 199: 1372–1381

Zhang Y, Liu T, Meyer CA, Eeckhoute J, Johnson DS, Bernstein BE, Nusbaum C, Myers RM, Brown M, Li W, et al (2008) Model-based Analysis of ChIP-Seq (MACS). 9: R137

